# Long non-coding RNA repertoire and regulation by nuclear exosome, cytoplasmic exonuclease and RNAi in fission yeast

**DOI:** 10.1101/158477

**Authors:** Sophie R Atkinson, Samuel Marguerat, Danny A Bitton, Maria Rodríguez-López, Charalampos Rallis, Jean-François Lemay, Cristina Cotobal, Michal Malecki, Juan Mata, François Bachand, Jürg Bähler

**Affiliations:** University College London, Research Department of Genetics, Evolution & Environment and UCL Cancer Institute, London, WC1E 6BT, UK.; MRC London Institute of Medical Sciences (LMS), London W12 0NN UK; Institute of Clinical Sciences (ICS), Faculty of Medicine, Imperial College London, London W12 0NN UK; Université de Sherbrooke, Department of Biochemistry, Sherbrooke, Quebec J1H 5N4, Canada.; Department of Biochemistry, University of Cambridge, Cambridge CB2 1QW, UK.; Current address: University of East London, School of Health Sports and Bioscience, London, EN15 4LZ, London, UK.

**Keywords:** pervasive transcription, NMD pathway, Schizosaccharomyces pombe, RNA degradation, antisense RNA

## Abstract

Transcriptomes feature pervasive, but poorly defined long non-coding RNAs (lncRNAs). We identify 5775 novel lncRNAs in *Schizosaccharomyces pombe*, nearly 4-times the previously annotated lncRNAs. Most lncRNAs become derepressed under genetic and physiological perturbations, especially during late meiosis. These lncRNAs are targeted by three RNA-processing pathways: the nuclear exosome, cytoplasmic exonuclease and RNAi, with substantial coordination and redundancy among pathways. We classify lncRNAs into cryptic unstable transcripts (CUTs), Xrn1-sensitive unstable transcripts (XUTs), and Dicer-sensitive unstable transcripts (DUTs). XUTs and DUTs are enriched for antisense lncRNAs, while CUTs are often bidirectional and actively translated. The cytoplasmic exonuclease and RNAi repress thousands of meiotically induced RNAs. Antisense lncRNA and sense mRNA expression often negatively correlate in the physiological, but not the genetic conditions. Intergenic and bidirectional lncRNAs emerge from nucleosome-depleted regions, upstream of positioned nucleosomes. This broad survey of the *S. pombe* lncRNA repertoire and characteristics provides a rich resource for functional analyses.

## Introduction

Genomes are more pervasively transcribed than expected from their protein-coding sequences. For example, 70-80% of the human genome is transcribed but less than 2% encodes proteins (Djebali et al. 2012). So-called long non-coding RNAs (lncRNAs), which exceed 200 nucleotides in length but lack long open reading frames, make up a substantial and diverse portion of the non-coding transcriptome. The functions, if any, of most lncRNAs are not known, although several have well-defined roles in gene regulation and other cellular processes, and are also implicated in diseases (Geisler and Coller 2013; Batista and Chang 2013; Mercer and Mattick 2013; Guttman and Rinn 2012; Rinn and Chang 2012). The lncRNAs are often lowly expressed but show more changes in expression levels between different tissues or conditions than do the protein-coding messenger RNAs (mRNAs) (Cabili et al. 2011; Derrien et al. 2012; Pauli et al. 2012; Hon et al. 2017). In general, lncRNAs are transcribed by RNA polymerase II and seem to be capped and polyadenylated (Cabili et al. 2011; Derrien et al. 2012; Guttman et al. 2009), although the patterns of transcription and RNA processing can radically differ between mRNAs and lncRNAs (Schlackow et al. 2017; Tuck and Tollervey 2013; Quinn and Chang 2016; Mukherjee et al. 2017). Some lncRNAs engage with ribosomes, which can trigger nonsense-mediated decay (NMD) to dampen their expression but may also produce functional peptides in some cases (de Andres-Pablo et al. 2017; Quinn and Chang 2016; Malabat et al. 2015; Wery et al. 2016).

Given the profusion, diversity and low expression of lncRNAs, their full description and annotation is still ongoing and evolving (Mattick and Rinn 2015; St. Laurent et al. 2015; Atkinson et al. 2012). A conceptually simple way to classify lncRNA genes is by their position relative to neighbouring coding genes. For example, long intervening non-coding RNAs (lincRNAs), transcribed from intergenic regions that do not overlap any mRNAs, have been the subject of much research in mammalian cells (Ulitsky and Bartel 2013; Rinn and Chang 2012; Schlackow et al. 2017). Antisense lncRNAs are transcribed in the opposite direction to mRNAs with which they completely or partially overlap; they can affect sense transcript levels via diverse mechanisms (Pelechano and Steinmetz 2013; Mellor et al. 2016). Bidirectional lncRNAs, on the other hand, emerge close to the transcriptional start site of coding genes but run in the opposite direction; most eukaryotic promoters initiate divergent transcription leading to widespread bidirectional lncRNAs, although transcriptional elongation is often only productive in the sense direction (Quinn and Chang 2016; Grzechnik et al. 2014). Bidirectional transcription has been proposed to drive the origination of new genes (Wu and Sharp 2013) and modulate gene-expression noise (Wang et al. 2011).

Pervasive transcription is potentially harmful as it can affect the expression of coding genes (Mellor et al. 2016), and nascent RNAs can compromise genome stability (Li and Manley 2006). Cells therefore apply RNA surveillance systems to keep the expression of lncRNAs in check (Jensen et al. 2013). Many lncRNAs are actively degraded shortly after transcription, suggesting that they could reflect transcriptional noise, byproducts of coding transcription, or that the act of transcription, rather than the lncRNA itself, is functionally important (Jensen et al. 2013). Different RNA-decay pathways preferentially target distinct sets of lncRNAs, which has been used for lncRNA classification in budding yeast. The ‘cryptic unstable transcripts’ (CUTs) accumulate in cells lacking Rrp6 (Xu et al. 2009; Neil et al. 2009), a nuclear-specific catalytic subunit of the RNA exosome (Houseley and Tollervey 2009; Kilchert et al. 2016). Conversely, the ‘stable unannotated transcripts’ (SUTs) are less affected by Rrp6 (Xu et al. 2009; Neil et al. 2009). The human ‘promoter upstream transcripts’ (PROMPTs) are analogous to CUTs (Preker et al. 2008). The ‘Xrn1-sensitive unstable transcripts’ (XUTs) accumulate in cells lacking the cytoplasmic 5’ exonuclease Xrn1 and are often antisense to mRNAs (van Dijk et al. 2011; Houseley and Tollervey 2009). ‘Nrd1-unterminated transcripts’ (NUTs) and ‘Reb1p-dependent unstable transcripts’ (RUTs) are controlled via different mechanisms of transcriptional termination linked to RNA degradation (Schulz et al. 2013; Colin et al. 2014). These classes, while useful, are somewhat arbitrary and overlapping as most lncRNAs can be targeted by different pathways, especially if one pathway is absent in mutant cells (Jensen et al. 2013).

The fission yeast *Schizosaccharomyces pombe* provides a potent complementary model system to study gene regulation. In some respects, RNA metabolism of fission yeast is more similar to metazoan cells than budding yeast. For example, RNA interference (RNAi) (Castel and Martienssen 2013), RNA uridylation (Schmidt et al. 2011), and Pab2/PABPN1-dependent RNA degradation (Lemay et al. 2010; Lemieux et al. 2011; Beaulieu et al. 2012) are conserved from fission yeast to humans, but absent in budding yeast. Several genome-wide studies have uncovered widespread lncRNAs (Dutrow et al. 2008; Wilhelm et al. 2008; Rhind et al. 2011; Eser et al. 2016), and over 1500 lncRNAs are currently annotated in the PomBase model organism database (McDowall et al. 2014). Studies with natural isolates of *S. pombe* revealed that only the most highly expressed lncRNAs show purifying selection (Jeffares et al. 2015), but the regulation of many lncRNAs is affected by expression quantitative trait loci (Clément-Ziza et al. 2014). Over 85% of the annotated *S. pombe* lncRNAs are expressed below one copy per cell during proliferation, and over 97% appear to be polyadenylated (Marguerat et al. 2012). Ribosome profiling showed that as many as 24% of lncRNAs are actively translated (Duncan and Mata 2014). As in other organisms, a large proportion of the *S. pombe* lncRNAs are antisense to mRNAs and have been implicated in controlling the meiotic gene expression programme (Ni et al. 2010; Chen et al. 2012; Bitton et al. 2011). Diverse chromatin factors function in suppressing antisense and other lncRNAs in *S. pombe*, including the HIRA histone chaperone (Anderson et al. 2009), the histone variant H2A.Z (Zofall et al. 2009; Clément-Ziza et al. 2014), the Clr4/Suv39 methyltransferase together with RNAi (Zhang et al. 2011), the Spt6 histone chaperone (DeGennaro et al. 2013), and the CHD1 chromatin remodeller (Pointner et al. 2012; Hennig et al. 2012; Shim et al. 2012). Several lncRNAs have been functionally characterized in *S. pombe*: *meiRNA* controls meiotic differentiation and chromosome pairing (Yamashita et al. 2016; Ding et al. 2012), stress-induced lncRNAs activate expression of the downstream *fbp1* gene during glucose starvation (Hirota et al. 2008; Oda et al. 2015), *adh1AS* is an antisense inhibitor of the *adh1* gene during zinc limitation (Ehrensberger et al. 2013), *prt* recruits the exosome to control phosphate-tuned *pho1* expression (Shah et al. 2014; Ard et al. 2014), *nc-tgp1* inhibits the phosphate-responsive permease *tgp1* gene by transcriptional interference (Ard et al. 2014), and *SPNCRNA.1164* regulates expression of the *atf1* transcription-factor gene in *trans* during oxidative stress (Leong et al. 2014). Naturally, these studies only scratch the surface, and the non-coding transcriptome and any of its functions remain relatively poorly defined in fission yeast and other organisms.

Transcriptome analyses under selective conditions, such as in RNA-processing mutants, have proven useful to define lncRNAs in budding yeast. Here we analyse transcriptome sequencing under multiple genetic and physiological perturbations in fission yeast to maximize the detection and initial characterization of lncRNAs. Some of these RNA-seq samples have previously been analyzed with respect to mRNA processing and expression (Bitton et al. 2015a; Lemieux et al. 2011; Schlackow et al. 2013; Marguerat et al. 2012; Lemay et al. 2014). They interrogate pathways, such as RNAi and Pab2/PABPN1, which are conserved in humans but not in budding yeast. We identify 5775 novel, unannotated lncRNAs, in addition to the previously annotated lncRNAs. The expression of lncRNAs is more extensively regulated in stationary phase, quiescence and, most notably, meiotic differentiation than the expression of mRNAs. Many lncRNAs comprise unstable transcripts that are degraded by three partially overlapping RNA-processing pathways. Analogous to budding yeast, we classify the unstable lncRNAs targeted by Rrp6, Dcr1 and Exo2 into CUTs, DUTs and XUTs, respectively. We further analyse the positions and expression of all novel and annotated lncRNAs with respect to neighbouring mRNAs, and other biological characteristics such as translation and nucleosome patterns. This extensive study provides a framework for functional characterization of lncRNAs in fission yeast and beyond.

## Results

### Detection of novel lncRNAs

To broadly identify lncRNAs in fission yeast, we examined strand-specific RNA-seq data acquired under multiple genetic and physiological conditions. We analyzed the transcriptomes of twelve RNA-processing mutants to facilitate detection of RNAs that may be rapidly degraded (***Supplementary file 1***). This mutant panel affects proteins for key pathways of RNA processing and degradation: Rrp6, a 3’-5’ exonuclease of the nuclear RNA exosome (Harigaya et al. 2006; Lemay et al. 2014); Dis3, a 3’-5’ exonuclease/endonuclease of the core RNA exosome (Wang et al. 2008); Ago1 (Argonaute), Dcr1 (RNase III-like Dicer) and Rdp1 (RNA-dependent RNA polymerase) of the RNAi pathway (Volpe et al. 2002); Exo2, a cytoplasmic 5’ exonuclease (ortholog of XRN1) (Houseley and Tollervey 2009); Ski7, a cytoplasmic cofactor which links the Ski complex to the exosome (Lemay et al. 2010); Cid14, a polyA polymerase of the TRAMP complex which is a cofactor of the nuclear RNA exosome (Wang et al. 2008); Pab2, a polyA-binding protein targeting RNAs to the nuclear exosome (PABPN1 ortholog) (Lemieux et al. 2011); Pan2, a deadenylase of the Pan2–Pan3 complex (Wolf and Passmore 2014); and Upf1, an ATP-dependent RNA helicase of the NMD pathway (Rodríguez-Gabriel et al. 2006). We also analyzed transcriptomes under nine physiological conditions to sample key cellular states (***Supplementary file 2***): two timepoints of stationary phase after glucose depletion (100 and 50% cell viability); two timepoints of quiescence/cellular ageing (days 1 and 7 after nitrogen removal); and five timepoints of meiotic differention (0 to 8 h after triggering meiosis).

To detect novel lncRNAs longer than 200 nucleotides, we designed a segmentation heuristic, optimised by its ability to detect the previously annotated lncRNAs (Materials and methods). Applying this approach to the RNA-seq data covering the genetic and physiological perturbations, we identified 5775 novel, unannotated lncRNAs in addition to the ~1550 previously annotated lncRNAs. We assigned systematic names, *SPNCRNA.2000* to *SPNCRNA.7774*, to these novel lncRNAs (***Supplementary file 3***). Of these unannotated lncRNAs, 159 fully or partially overlapped on the same strand with 171 of the 487 novel lncRNAs recently reported (Eser et al. 2016) (***Supplementary file 4***). The novel lncRNAs were generally shorter (mean 797 nt) than the annotated lncRNAs (mean 1233 nt) and mRNAs (mean 2148 nt). While some sequence library protocols can generate spurious antisense RNAs (Perocchi et al. 2007), our protocol is resilient to this artefact as it relies on ligating two RNA oligonucleotides to fragmented mRNAs. We mostly analyzed polyA-enriched samples, because almost all *S. pombe* lncRNAs are polyadenylated (Marguerat et al. 2012). As a control, we compared the data to samples depleted for ribosomal RNA (rRNA) in *rrp6Δ* and *exo2Δ* mutants, which confirmed that the great majority of lncRNAs are polyadenylated (*Figure 1-figure supplement 1*).

None of the novel lncRNAs overlapped with mRNAs on the same strand using current PomBase annotations, and only 21 overlapped using CAGE (Li et al. 2015) and polyA data (Mata 2013), respectively, for transcription start and termination sites (***Supplementary file 5***). Thus, the novel lncRNAs do not represent alternative transcription start or termination sites of known mRNAs. On the other hand, 3650 of 5138 protein-coding regions (71%) overlapped by at least 10 nucleotides with antisense lncRNAs, either annotated or novel. The 1461 (28.4%) of coding regions not associated with antisense lncRNAs were enriched for the 20% shortest mRNAs (*p* ~1.6e^−16^), including those encoding ribosomal proteins (*p* ~1.1e^−5^), as well as for several features associated with high gene expression, including high mRNA levels (*p* ~0.004) (Pancaldi et al. 2010), stable mRNAs (*p* ~2.1e^−9^) (Hasan et al. 2014), and mRNAs that show high RNA polymerase II occupancy (*p* ~2.4e^−5^) and high ribosome density (*p* ~1.2e^−41^) (Lackner et al. 2007). These enrichments suggest that highly expressed genes are either protected from antisense transcripts or interfere with antisense transcription.

*Figure 1* shows the relative expression changes of all mRNAs, annotated lncRNAs and novel lncRNAs in the genetic and physiological conditions. For all conditions, differential expression was determined relative to three reference samples (exponentially proliferating wild-type and auxotrophic control cells in minimal medium with supplements; ***Supplementary file 2***). This panel of reference samples included different genetic markers and supplements used for the other conditions. Only 403 RNAs, including 184 novel lncRNAs, were differentially expressed between wild-type and auxotrophic control cells grown in rich yeast extract (YE) medium compared to minimal media (>2-fold change, p <0.05). Thus, the growth medium and auxtrophic markers only minimally affected lncRNA expression. All differential expression data are available in ***Supplementary file 3***.

**Figure 1.**
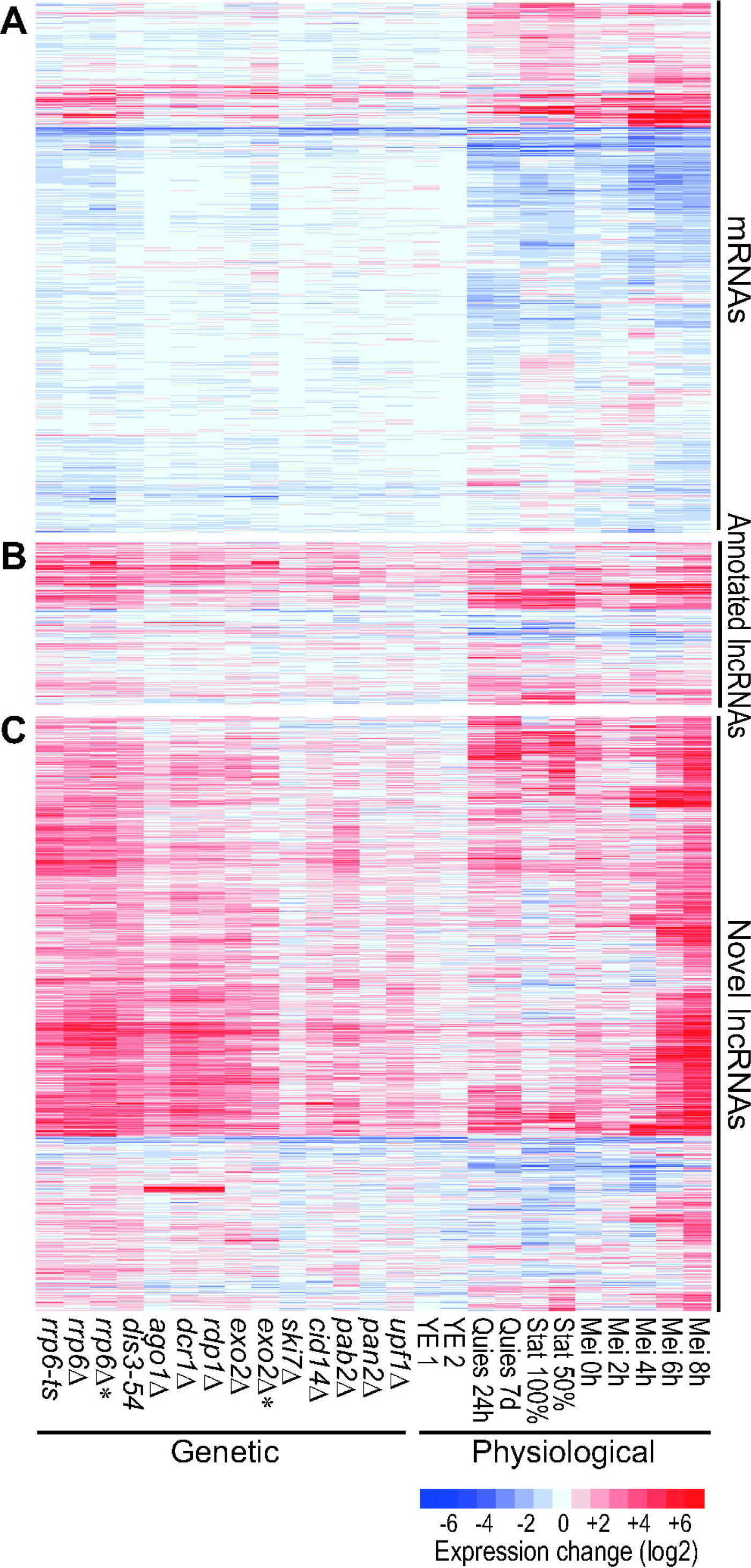
Hierarchical clustering of gene expression in different RNA-processing mutants and physiological conditions. Expression profiles are shown for (**A**) all 5177 mRNAs, (**B**) 1573 annotated lncRNAs, and (**C**) 5775 novel, unannotated lncRNAs. Changes in RNA levels in response to the different genetic and physiological conditions (indicated at bottom) relative to control cells grown in minimal medium are color-coded as shown in the color-legend at bottom right (log2 expression ratios). The *rrp6* and *exo2* samples indicated by asterisks has been depleted for rRNA instead of polyA purification used for all other samples. Data obtained from these two types of RNA-seq libraries are compared in *Figure 1-figure supplement 1*. Details on strains and conditions are provided in ***Supplementary files 1 and 2***, and all expression data are provided in ***Supplementary file 3***.

About 50% of the mRNAs were differentially expressed, both induced and repressed, in the different physiological conditions, but much less so in the different genetic conditions (*Figure 1A*, *Figure 2*). The mRNAs up-regulated in the exosome, *pab2*, and RNAi mutants were enriched for Gene Ontology (GO) terms related to meiosis, consistent with reported roles of the corresponding proteins in meiotic gene silencing (St-André et al. 2010; Yamanaka et al. 2010, 2013). Notably, mRNAs up-regulated in the *exo2* mutant were strongly and specifically enriched for middle meiotic genes (Mata et al. 2002).

**Figure 2.**
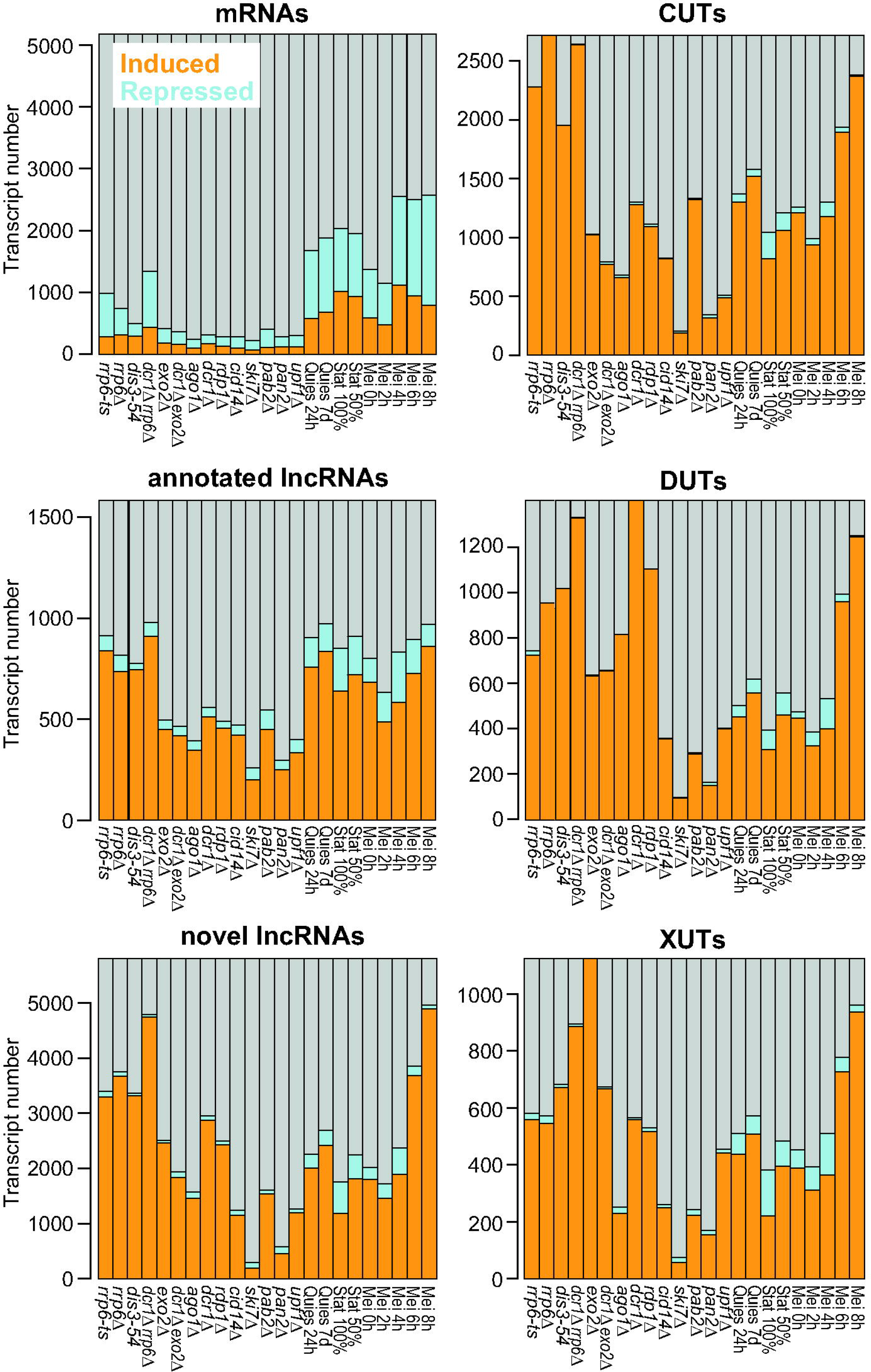
Histograms showing numbers and proportions of induced (orange), repressed (blue) and all other (grey) transcripts for the different RNA classes as indicated. Differentially expressed genes were defined as those being ≥2-fold induced (mean of two biological repeats) or repressed and showing significant changes (p <0.05) compared to reference as determined by DESeq2.

*Figures 1B and C* show the expression of the previously annotated and novel lncRNAs, respectively. Compared to mRNAs, much higher proportions and numbers of lncRNAs featured strong differential expression, mostly induced, under the different genetic and physiological conditions (*Figure 1B,C*, *Figure 2*). The following mutants led to the most pronounced effects on lncRNA expression: nuclear exosome (*rrp6-ts, rrp6Δ, dis3-54*), RNAi (*ago1Δ, dcr1Δ, rdp1Δ*), and the cytoplasmic exonuclease (*exo2Δ*). In nuclear exosome mutants, most lncRNAs were strongly derepressed, in stark contrast to the mRNAs which showed a higher proportion of repressed transcripts in this condition (*Figure 2, Figure 3A*). Many of the novel lncRNAs in particular were also strongly derepressed in RNAi and *exo2* mutants and in meiotic cells (*Figure 3A*), most notably during late meiotic stages (Mei 6-8h; *Figure 1C*, *Figure 2*). On the other hand, the novel lncRNAs showed generally lower expression levels in both genetic and physiological conditions than the annotated lncRNAs, and much lower than the mRNAs (*Figure 3B*). Remarkably, only 54 novel lncRNAs were neither up- nor down regulated in any of the conditions (>2-fold change, p <0.05). This result argues against sequencing artefacts and supports that the levels of these transcripts are modulated in response to different biological conditions. Together, their low expression levels and induction under specialized conditions can explain why the novel lncRNAs have not been identified in previous studies (Wilhelm et al. 2008; Rhind et al. 2011; Eser et al. 2016).

**Figure 3.**
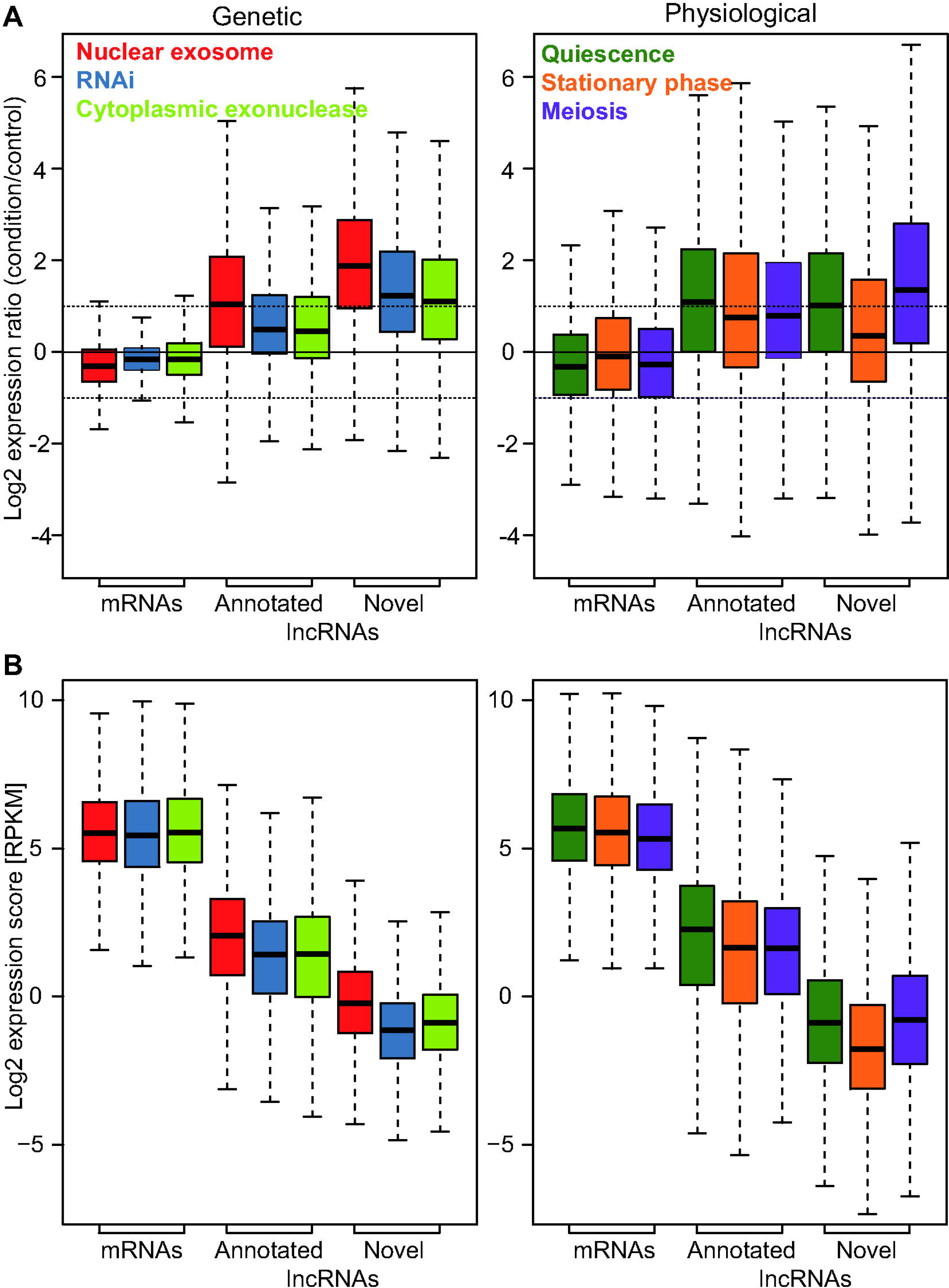
Gene expression in major groups of genetic and environmental conditions. (**A**) Left graph: box plot of expression ratios (condition relative to control) of all mRNAs, annotated and novel lncRNAs in nuclear exosome (*rrp6Δ, rrp6-ts*), RNAi (*ago1Δ, dcr1Δ, rdp1Δ*) and cytoplasmic exonuclease (*exo2Δ*) mutants. Right graph: as left but for quiescence, stationary phase and meiosis conditions. The horizontal dashed lines indicate 2-fold induction and repression.(**B**) As in panel **A**, but for expression levels (RPKM scores). All expression data are provided in ***Supplementary file 3***.

In conclusion, these findings indicate that the novel lncRNAs comprise many cryptic transcripts that are degraded by three main RNA-processing pathways during mitotic proliferation: the nuclear exosome, the RNAi machinery, and the cytoplasmic exonuclease Exo2. The other RNA processing factors analyzed here seem to play only minor or redundant roles in lncRNA regulation. Many lncRNAs become induced under specific physiological conditions when they might play specialized roles.

### Classification of lncRNAs into CUTs, DUTs and XUTs

In budding yeast, different groups of lncRNAs have been named according to the RNA-processing pathways controlling their expression. For example, CUTs are targeted for degradation by the nuclear exosome (Neil et al. 2009; Xu et al. 2009), and XUTs are targeted by the cytoplasmic exonuclease Xrn1 (ortholog of *S. pombe* Exo2) (van Dijk et al. 2011). We introduce an analogous classification of lncRNAs to provide a framework for analysis. Based on the panel of RNA-processing mutants tested here, the nuclear exosome, the cytoplasmic exonuclease, and the RNAi machinery are the three main pathways targeting lncRNAs (*Figure 1, Figure 3*). These three pathways thus provide a natural way to classify the lncRNAs as CUTs and XUTs (corresponding to the budding yeast classes of the same names) and DUTs (Dicer-sensitive unstable transcripts). DUTs define a novel class not applicable to budding yeast which lacks the RNAi machinery.

Using a fuzzy clustering approach (Materials and methods), we classified both the novel and previously annotated lncRNAs that were significantly derepressed in at least one of the following mutants: *rrp6Δ* (CUTs), *dcr1Δ* (DUTs) or *exo2Δ* (XUTs). Of 7308 lncRNAs, 2068 remained unclassified because they were not significantly derepressed in any of the three mutants (1896 lncRNAs) or could not be assigned to a single class (172 lncRNAs).

The remaining lncRNAs were classified into 2732 CUTs (493 annotated, 2239 novel), 1116 XUTs (181 annotated, 935 novel), and 1392 DUTs (209 annotated, 1183 novel). The resulting three classes were quite distinct (*Figure 4A*). These class associations are provided in ***Supplementary files 3 and 5***. As expected, the lncRNAs of a given class were most highly expressed on average in the mutant used to define the class, although they were also more highly expressed in the other two mutants than in wild-type cells (*Figure 4B*). This pattern was also evident when clustering the three lncRNA classes separately based on the expression changes in the different RNA-processing mutants: lncRNAs of a given class were most highly derepressed in the mutant used to define this class, and in mutants affecting the same pathway, but they also tended to be derepressed in mutants of other pathways (*Figure 4C*). These results show that the lncRNAs of a given class are not exclusively derepressed in the mutant used to define this class. Thus, lncRNAs can be degraded by different pathways, although one pathway is typically dominant for a given lncRNA which is used here for classification.

**Figure 4.**
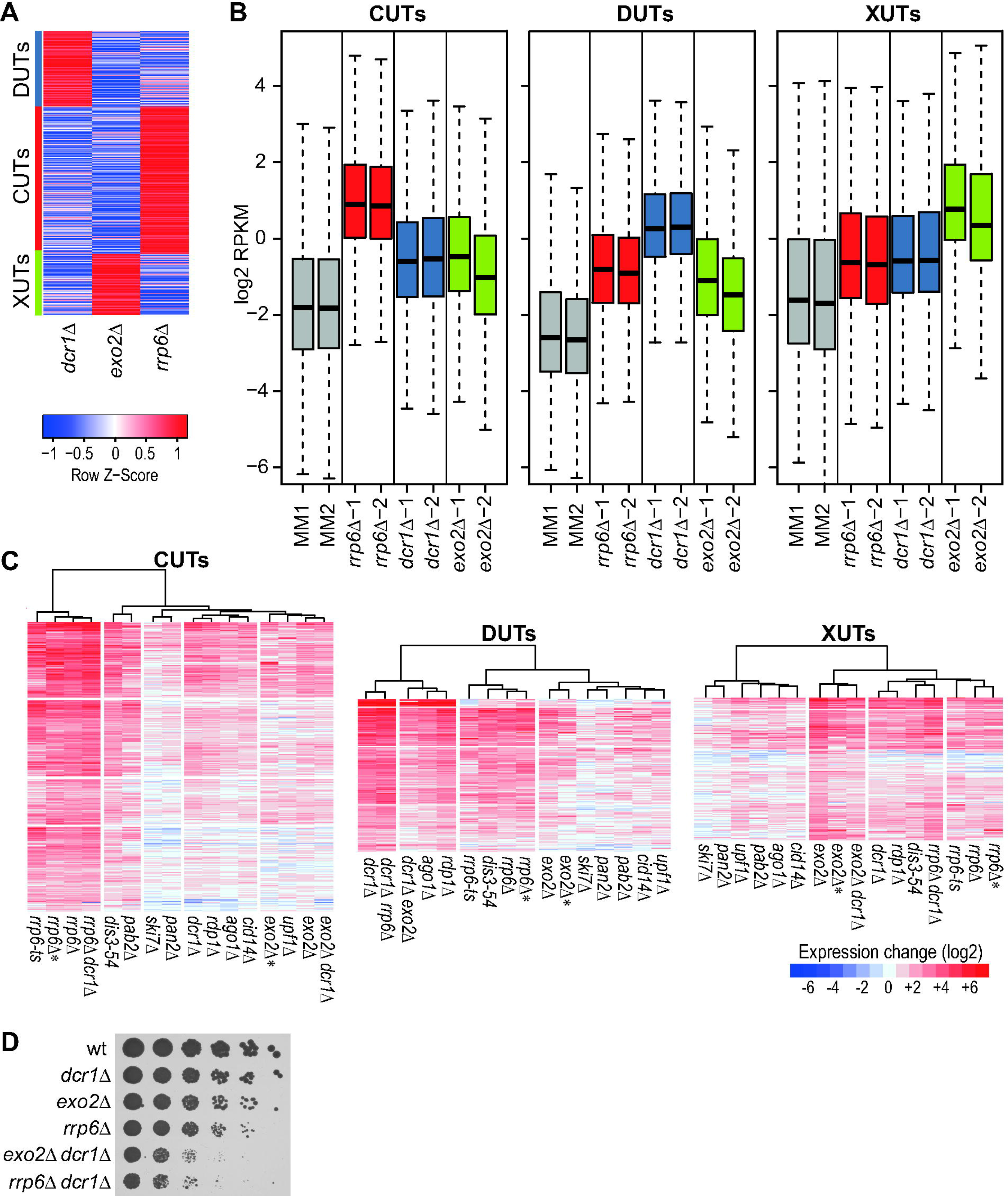
Classification of lncRNAs into CUTs, DUTs and XUTs. (**A**) The lncRNAs significantly induced (DESeq2; adjusted p ≤0.05) in *rrp6Δ*, *dcr1Δ* or *exo2Δ* mutants were clustered into CUTs, DUTs and XUTs, respectively, using the Mfuzz R package (default parameters, 3 clusters specified). The clustering shows the 5586 uniquely classified lncRNAs after filtering those with a membership score <0.7. The red/blue colours indicate the mean RPKM values in the 3 mutants as indicated, scaled by subtracting the mean of the row and division by the standard deviation of the row (z-score). The assigned clusters are indicated at left.(**B**) Box plots of RPKM values (log2) of all CUTs (left), DUTs (middle) and XUTs (right) in control (MM) and mutant cells as indicated. The data for the biological repeats 1 and 2 are ploted separately.(**C**) Hierarchical clustering of genetic conditions for all CUTs, DUTs and XUTs as indicated. Changes in RNA levels in response to the different genetic conditions (indicated at bottom) relative to control cells grown in minimal medium are color-coded as shown in the legend at bottom right (log2 expression ratios). All expression and classification data are provided in ***Supplementary file 3***.(**D**) Growth phenotypes of mutant cells. Six 5-fold serial dilutions of wild-type (wt) and mutant cells as indicated were spotted on YE medium plates and photographed after 3 days of growth at 32°C.

### Relationships between RNA-processing pathways targeting lncRNAs

To further dissect the functional relationships among the nuclear exosome, RNAi, and cytoplasmic exonuclease pathways, we attempted to construct double and triple mutants of *rrp6Δ, dcr1Δ* and *exo2Δ*. We could not obtain an *rrp6Δ exo2Δ* mutant from 24 tetrads dissected from the corresponding cross. Among the tetrads analyzed with three viable spores, the non-surviving spore was always of the *rrp6Δ exo2Δ* genotype, with a significant difference between observed and expected frequencies of wild-type, single and double mutant spores (chi-square test, p ~10^−4^; based on 61% surviving spores). We conclude that the *rrp6Δ exo2Δ* double mutant is not viable. This synthetic lethality indicates that the nuclear exosome and cytoplasmic exonuclease together exert an essential role.

Conversely, the *exo2Δ dcr1Δ* and *rrp6Δ dcr1Δ* double mutants were viable, although the latter showed stronger growth defects than either single mutant (*Figure 4D*). While the *exo2Δ dcr1Δ* cells were elongated like the *exo2Δ* cells (Szankasi and Smith 1996), the *rrp6Δ* and *rrp6Δ dcr1Δ* cells were of normal length, with the double mutant looking more irregular and sick (data not shown). These findings suggest that the nuclear exosome and RNAi machineries can back each other up to some extend. We also attempted to construct an *rrp6Δ exo2Δ dcr1Δ* triple mutant by mating of the *exo2Δ dcr1Δ* and *rrp6Δ dcr1Δ* double mutants. This mating only produced ~17% viable spores, suggesting that the RNAi machinery is required for spore survival. As expected, we did not obtain any triple mutant among 24 tetrads dissected from this cross (chi-square test, p ~10^−4^; based on 17% surviving spores). Thus, both the *rrp6Δ exo2Δ* double mutant and the *rrp6Δ exo2Δ dcr1Δ* triple mutant are not viable. Consistent with lncRNAs being targeted by multiple pathways (*Figure 4B,C*), these results point to some redundancy in function between the different RNA degradation pathways (Discussion).

We analyzed the transcriptomes of the *exo2Δ dcr1Δ* and *rrp6Δ dcr1Δ* double mutants by RNA-seq. The *rrp6Δ dcr1Δ* double mutant showed a greater number of derepressed lncRNAs than either single mutant, especially among the novel lncRNAs (*Figure 2; Figure 4C*). The 5647 lncRNAs that were significantly derepressed in the *rrp6Δ dcr1Δ* double mutant (expression ratio >2, p <0.05) included 2653 CUTs and 1317 DUTs, but also 884 XUTs and 793 unclassified lncRNAs. This result again shows that the nuclear exosome and RNAi have partially redundant roles and can back each other up with respect to many RNA targets. Moreover, these two nuclear pathways can also degrade most XUTs that are further targeted by the cytoplasmic exonuclease. These findings highlight a prominent role of the joint activity of the nuclear exosome and RNAi to suppress a large number of lncRNAs.

In contrast, the *exo2Δ dcr1Δ* double mutant showed fewer derepressed XUTs and DUTs than either single mutant (*Figure 2, Figure 4C*). The 2272 lncRNAs that were significantly derepressed in the double mutant included only 675 XUTs and 658 DUTs, but 786 CUTs and 153 unclassified lncRNAs. Taken together, the complex findings from single and double mutants indicate that the three RNA-processing pathways can all target a wide range of lncRNAs, with differential preference for some targets, and channeling of lncRNAs to alternative pathways in the mutants. The higher numbers of derepressed lncRNAs in *rrp6Δ* and *dcr1Δ* mutants (*Figure 2, Figure 4C*) indicate that most lncRNAs are degraded in the nucleus, most notably by the nuclear exosome.

### Classification of lncRNAs by neighbouring mRNA positions

We also classified all known and novel lncRNAs into the main types based on their positions relative to the neighbouring mRNAs: Bidirectional, Antisense and Intergenic (*Figure 5A*). The criteria for these assignments, and overlaps between different classes, are specified in Materials and methods. In total, we defined 1577 Bidirectional (539 annotated, 1038 novel), 4474 Antisense (575 annotated, 3899 novel) and 1189 Intergenic (356 annotated, 833 novel) lncRNAs, besides 108 that overlapped mRNAs in sense direction (103 annotated, 5 novel). Data for these lncRNA classes are provided in ***Supplementary file 5***. The Bidirectional lncRNAs were enriched for CUTs (*Figure 5A*). The Antisense lncRNAs were enriched for XUTs and, most notably, for DUTs, while the Intergenic lncRNAs were enriched for other, not classified lncRNAs (*Figure 5A*). Similar trends were also evident when analysing the lncRNA types the other ways round (*Figure 5B*): while most CUTs, DUTs and XUTs were Antisense, only the DUTs and XUTs were enriched for Antisense lncRNAs, while the CUTs were enriched for Bidirectional lnsRNAs.

**Figure 5.**
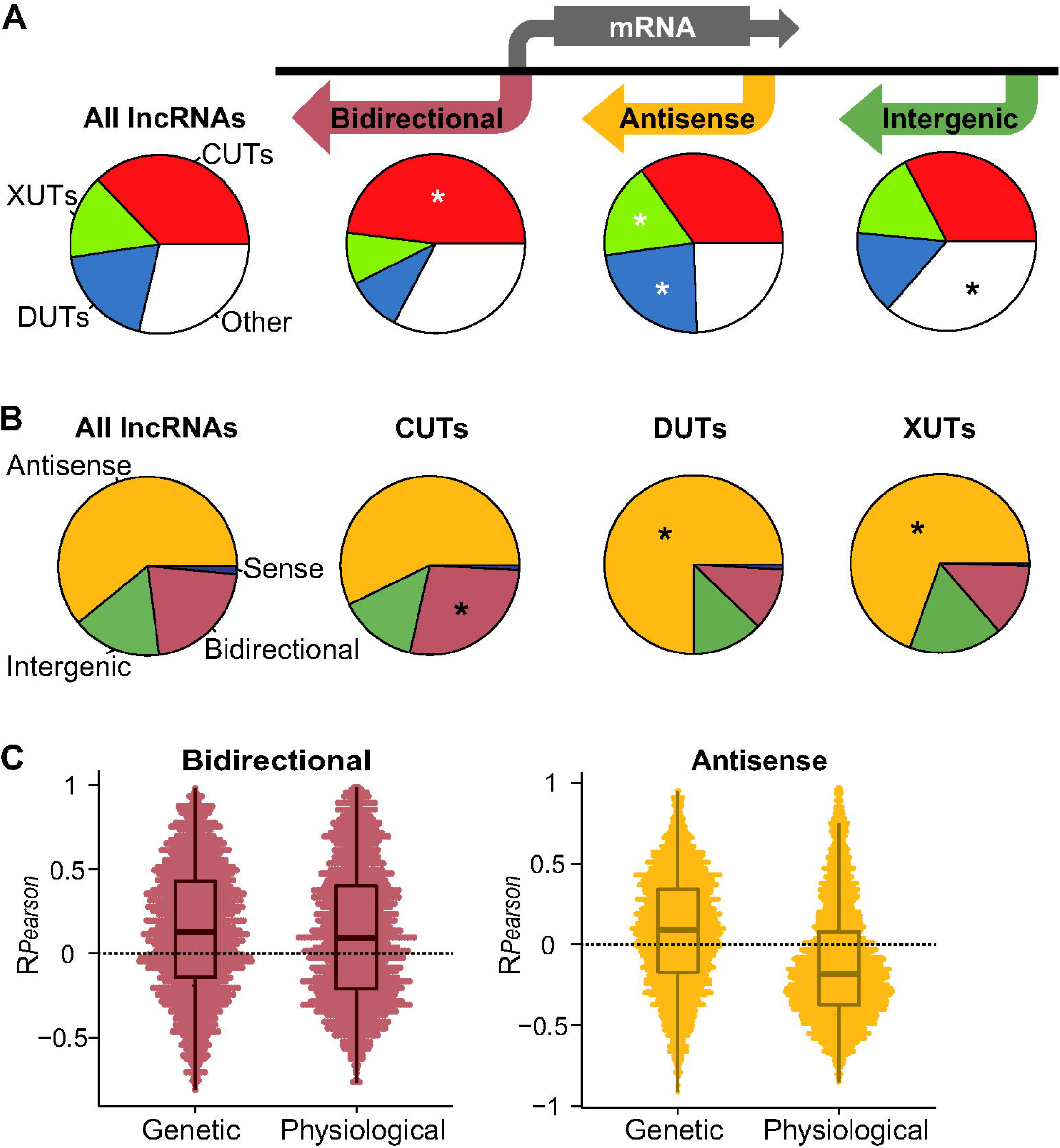
Analyses of lncRNAs by positions relative to mRNAs. (**A**) We grouped the annotated and novel lncRNAs into three main positional types as represented schematically: 1577 Bidirectional, 4474 Antisense and 1189 Intergenic RNAs, leaving only 108 lncRNAs that overlapped mRNAs in sense direction (Materials and methods). Pie charts of the corresponding proportions of CUTs, DUTs, XUTs and other lncRNAs are provided beneath each positional type, and also for all annotated and known lncRNAs. Significantly enriched slices are indicated with asterisks (*R prop.test function*, p <10^−6^).(**B**) Pie charts of the proportions of Bidirectional, Antisense, Intergenic and Sense lncRNAs for all (annotated and known) lncRNAs, and among the CUTs, DUTs and XUTs. Significantly enriched slices are indicated with asterisks (*R prop.test function*, p <10^−4^). (**C**) Pearson correlation coefficients for RPKM expression data of each Bidirectional lncRNA-mRNA pair (left) and each Antisense lncRNA-mRNA pair (right). The correlation data are shown separately for all genetic and physiological conditions as indicated on X-axes. For Antisense lncRNAs, the difference between the distributions in genetic *vs* physiological conditions is highly significant (P*Wilcoxon* = 4.6e^−170^). All classification data are provided in ***Supplementary file 5***.

### Expression patterns of lncRNAs

The expression of lncRNAs could be affected by the expression of neighboring mRNAs, or it could actively control the expression of neighbouring mRNAs. To analyze the relationship between Bidirectional and Antisense lncRNAs and their associated mRNAs, we calculated the correlation coefficients for expression levels of each lncRNA-mRNA pair across all genetic and physiological conditions. *Figure 5C* shows that the expression of the Bidirectional lncRNAs tended to positively correlate with the expression of their associated mRNAs, for both the genetic and physiological conditions. The Antisense lncRNAs, on the other hand, revealed substantial differences for genetic *vs* physiological conditions. They mainly correlated negatively with sense mRNA expression in the physiological, but not in genetic conditions (*Figure 5C*). Thus, accumulation of Antisense lncRNAs in the different RNA-processing mutants is generally not sufficient to repress mRNA levels. These striking contrasts between the two lncRNA classes and the different types of conditions provide clues about the lncRNA-mRNA expression relationships for Bidirectional and Antisense lncRNAs (Discussion).

The stabilisation of lncRNAs under specific physiological conditions (*Figure 1, Figure 3A*) could reflect specialized roles under these conditions. We checked whether the different lncRNAs showed distinct regulatory patterns by clustering the classes separately based on their expression changes in the different physiological conditions. When clustering CUTs, DUTs and XUTs, the physiological conditions always clustered into three main groups that showed distinct patterns of lncRNA expression (*Figure 6-figure supplement 1*): late meiosis, stationary phase (triggered by glucose limitation), and quiescence/early meiosis (both triggered by nitrogen limitation). These three groups reflect the major physiological states interrogated by the different conditions. The expression changes of CUTs, DUTs and XUTs, however, did not substantially differ across the physiological conditions. Most lncRNAs in all three classes, and among novel lncRNAs in particular, were strongly induced in late meiosis (Mei 8h; *Figure 2, Figure 6-figure supplement 1*). Many lncRNAs were also induced during stationary phase and quiescence/early meiosis, with CUTs being relatively most conspicuous in these conditions (*Figure 2, Figure 6-figure supplement 1*).

We also analyzed the lncRNAs classified by their positions relative to neighbouring mRNAs. As expected from *Figure 5A,B*, the nuclear exosome mutants showed the highest proportion of derepressed Bidirectional lncRNAs but were also prominent in derepressing Antisense and Intergenic lncRNAs (*Figure 6A*). The *rrp6Δ dcr1Δ* double mutant led to derepression of a large number of lncRNAs, particularly Intergenic and Antisense lncRNAs (*Figure 6A*). Among the physiological conditions, late meiosis (Mei 8h) showed the highest proportions of induced lncRNAs for all three classes; this response was most notable for Antisense lncRNAs, many of which were highly induced, while much fewer Intergenic lncRNAs were induced (*Figure 6A,B*). Overall, the Bidirectional, Antisense and Intergenic lncRNAs showed stronger class-specific expression signatures in the physiological conditions than did the CUTs, XUTs and DUTs (*Figure 6B vs Figure 6-figure supplement 1*). But expression patterns across the different physiological conditions were not sufficiently distinct to predict class membership based on these patterns.

**Figure 6.**
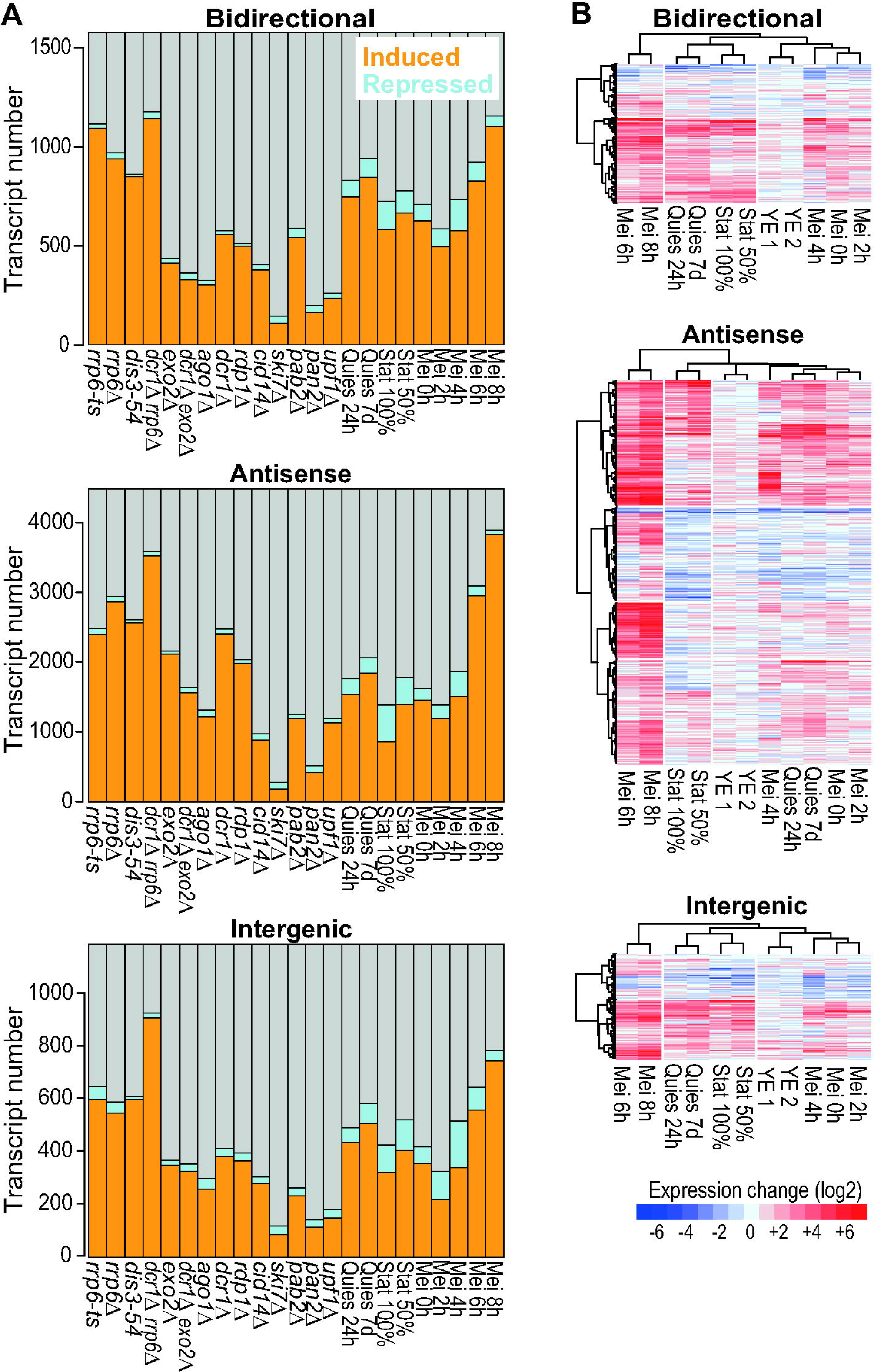
Expression patterns of Bidirectional, Antisense and Intergenic lncRNAs. (**A**) Histograms showing the numbers and proportions of the induced (orange), repressed (blue) and all other (grey) transcripts for the different lncRNA classes as indicated. Differentially expressed genes were defined as those being >2-fold induced (average of two biological repeats) or repressed and showing significant changes (p <0.05) compared to reference as determined by DESeq2.(**B**) Hierarchical clustering of physiological conditions for different lncRNA classes as indicated. Changes in RNA levels in response to the different physiological conditions (indicated at bottom) relative to control cells grown in minimal medium are color-coded as shown in the legend at bottom (log2 expression ratios). Hierarchical clustering of physiological conditions for CUTs, DUTs and XUTs is shown in *Figure 6-figure supplement 1*. All expression and classification data are provided in ***Supplementary files 3 and 5***.

### Nucleosome profiles of lncRNA regions

Transcribed regions are often accompanied by distinct patterns of nucleosome distribution. As a proxy for nucleosome distributions around lncRNA regions, we sequenced mononucleosomal DNA to compare the chromatin organization between protein-coding and non-coding transcribed regions in proliferating wild-type cells. We only analyzed lncRNA regions that did not overlap with any mRNA regions to minimize confounding data from coding transcription (although some mRNA transcription start sites may remain among the Bidirectional lncRNA data). *Figure 7* shows that both Bidirectional and Intergenic lncRNAs initiated in nucleosome-depleted regions, at the 5’-end of a positioned nucleosome. This feature was shared with mRNA regions, although these regions showed much higher nucleosome densities and a higher order of subsequent nucleosomes. In general, intergenic regions showed lower nucleosome densities than did mRNA regions, as has been observed before (Lantermann et al. 2010). We conclude that there are both similarities (around the transcription start site) and differences in chromatin organization between coding and non-coding regions.

**Figure 7.**
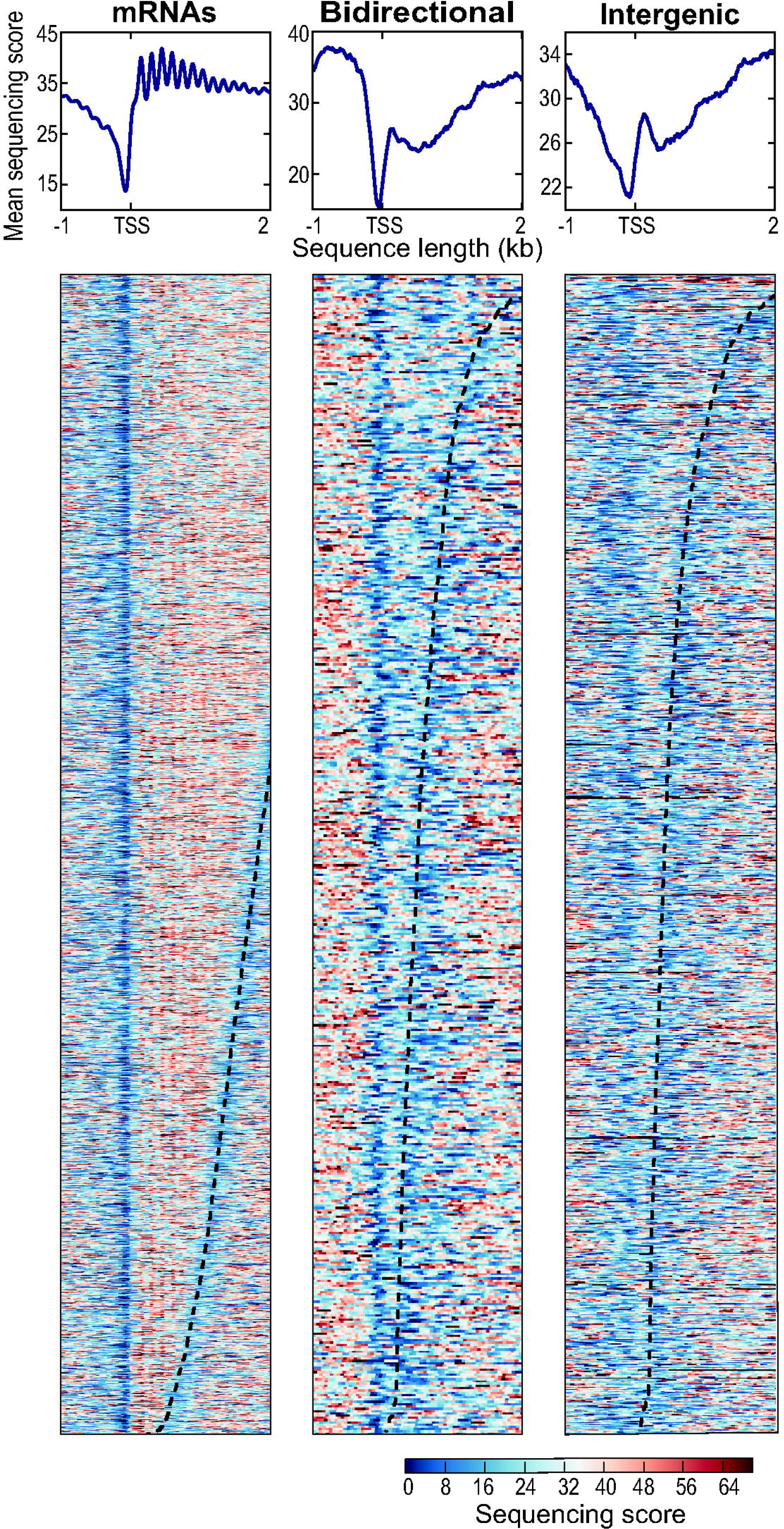
Nucleosome patterns for coding and non-coding transcribed regions. Nucleosome profiling data for all mRNA loci (left), 509 Bidirectional lncRNA loci and 1119 Intergenic lncRNA loci (right) in proliferating wild-type cells. Bidirectional lncRNA loci that overlap mRNAs in antisense direction were not included. Seventy Intergenic lncRNA loci that showed unusually high histone occupancies were omitted (***Supplementary file 8***). The top graphs show average nucleosome profiles for the different types of transcribed regions, aligned to the transcription start sites (TSS). The lower graphs show heatmaps for the first two kilobases of all transcribed regions analysed, ordered by transcript length from top (longest RNAs) to bottom (shortest RNAs). Sequencing scores are color-coded as shown in legend at bottom right.

### Translation of lncRNAs

Ribosome profiling before and during meiotic differentiation has revealed that as much as 24% of the annotated lncRNAs are actively translated (Duncan and Mata 2014). Such translation typically increases during meiosis and involves short open reading frames (ORFs), often more than one per lncRNA. In addition, translation has been detected in numerous unannotated regions of the genome (Duncan and Mata 2014). We analyzed the ribosome-profiling data from Duncan and Mata (2014), covering the whole genome of proliferating and meiotic cells, to assess translation of the different classes of annotated and novel lncRNAs defined above. Details of all 771 translated non-coding regions are provided in ***Supplementary file 6***.

Table 1 shows the numbers and percentages of actively translated lncRNAs, both for all translated regions of at least one codon and for a conservative set of translated regions with at least ten codons. Overall, the novel lncRNAs showed a much lower proportion of translated transcripts than the annotated lncRNAs. This result likely reflects their lower expression levels (*Figure 3B*). Such low expression makes it harder to obtain sufficient ribosome-profiling reads to determine translation for most lncRNAs, especially because no ribosome profiling data were available for most conditions in which the novel lncRNAs became derepressed. Nevertheless, 66 or 148 novel lncRNAs were found to be actively translated using the more or less conservative cutoff, respectively. Among the main classes, the Bidirectional lncRNAs showed the largest numbers and proportions of translated RNAs, which were highly enriched (*Table 1*; p = 1.7e^−26^). Moreover, up to 30% of the 108 previously annotated lncRNAs overlapping mRNAs in sense direction were actively translated. This finding suggests that some of these RNAs are alternative ‘untranslated regions’ of the mRNAs with translation of short upstream ORFs (Duncan and Mata 2014).

**Table 1.**
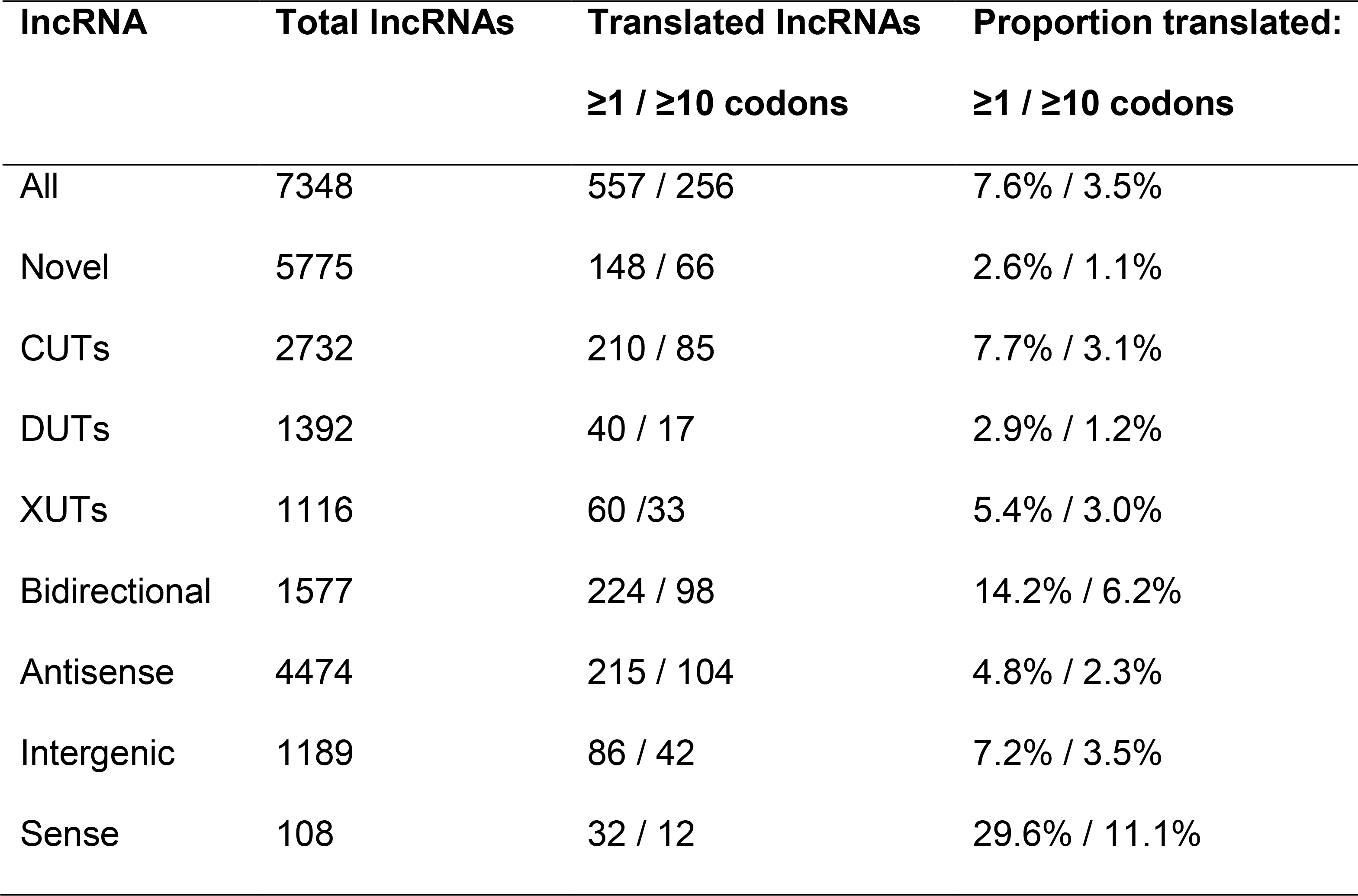
Data of actively translated lncRNA classes.

| lncRNA | Total lncRNAs | Translated lncRNAs | Proportion translated: |
| --- | --- | --- | --- |
|  |  | ≥1 / ≥10 codons | ≥1 / ≥10 codons |
| All | 7348 | 557 / 256 | 7.6% / 3.5% |
| Novel | 5775 | 148 / 66 | 2.6% / 1.1% |
| CUTs | 2732 | 210 / 85 | 7.7% / 3.1% |
| DUTs | 1392 | 40 / 17 | 2.9% / 1.2% |
| XUTs | 1116 | 60 / 33 | 5.4% / 3.0% |
| Bidirectional | 1577 | 224 / 98 | 14.2% / 6.2% |
| Antisense | 4474 | 215 / 104 | 4.8% / 2.3% |
| Intergenic | 1189 | 86 / 42 | 7.2% / 3.5% |
| Sense | 108 | 32 / 12 | 29.6% / 11.1% |

It is not clear to what extent any stable, functional peptides are generated by all this translational activity. The engagement of lncRNAs with ribosomes could trigger NMD and degradation via the cytoplasmic exonuclease (de Andres-Pablo et al. 2017; Quinn and Chang 2016; Malabat et al. 2015). We therefore checked whether the translated lncRNAs were enriched among those being derepressed in the *upf1Δ* mutant which is defective for NMD (Rodríguez-Gabriel et al. 2006). There were no significant overlaps between the translated RNAs and the novel, annotated or all lncRNAs derepressed in *upf1Δ* cells. Moreover, despite being enriched for translated lncRNAs, the Bidirectional lncRNAs showed no significant overlap with RNAs derepressed in *upf1Δ* cells. This finding is consistent with Bidirectional lncRNAs being mainly targeted by the nuclear exosome rather than the cytoplasmic exonuclease (*Figure 5A,B*). Conversely, the translated XUTs and Antisense lncRNAs were both significantly enriched for RNAs derepressed in *upf1Δ* cells (p = 6.7e^−24^ and 1.5e^−25^, respectively). This result is consistent with the cytoplasmic exonuclease being the major pathway for NMD-mediated RNA degradation. Together, these findings suggest that engaging with ribosomes triggers the degradation of many XUTs and Antisense lncRNAs, but not of Bidirectional lncRNAs which are thus more likely to produce peptides.

## Discussion

This study has uncovered 5775 novel lncRNAs. Compared to mRNAs and previously annotated lncRNAs, these novel lncRNAs are subject to stronger and more widespread differential expression, mostly induction, in response to multiple genetic and physiological conditions. Analysis of lncRNA expression across a broad panel of RNA-processing mutants indicates that the nuclear exosome, the RNAi pathway and the cytoplasmic exonuclease are key pathways targeting lncRNAs. Analogous to budding yeast, we have classified lncRNAs into CUTs, DUTs and XUTs, defined by the pathway which preferentially degrades them. Notably, mRNAs are much less affected by the absence of these RNA-processing pathways than are lncRNAs (*Figures 1-3*), suggesting that lncRNAs are important targets of these pathways. Unstable lncRNAs have been most extensively described at a genome-wide level in budding yeast which guided our classification. The lncRNA classes defined here show both similarities and differences to the classes defined in budding yeast, as highlighted below.

The ~2000 budding yeast CUTs are derepressed upon deletion of the nuclear-specific exosome subunit Rrp6 and transcribed divergently from mRNAs with which they positively correlate in expression (Neil et al. 2009; Xu et al. 2009). These findings are similar to our results (*Figure 4, Figure 5*). A difference, however, is that budding yeast CUTs are greatly stabilized by loss of Trf4, a key component of the exosome-targeting TRAMP complex (Wlotzka et al. 2011; Frenk et al. 2014), while fission yeast CUTs are only marginally affected by loss of the TRAMP subunit Cid14 (*Figure 1*; *Figure 2*) and Mtr4 (data not shown). This result indicates a TRAMP-independent mechanism of exosomal degradation of CUTs in fission yeast. Indeed, the polyA-binding protein Pab2, functioning in a complex called MTREC, targets meiotic or unspliced mRNAs and lncRNAs in fission yeast (Yamanaka et al. 2010; McPheeters et al. 2009; Lee et al. 2013; Egan et al. 2014; Zhou et al. 2015; St-André et al. 2010). Our data support a Pab2-dependent mechanism for targeting CUTs, showing a stronger derepression of CUTs in *pab2* mutants than in *cid14* mutants, although both mutants show much smaller effects than the exosome mutants *rrp6* and *dis3* (*Figures 1-3*). Also in human cells, many lncRNAs are targeted for exosomal degradation by PABPN1, an ortholg of Pab2 (Beaulieu et al. 2012; Meola et al. 2016). Reminiscent of yeast CUTs, exosome depletion in mammals has revealed lncRNAs divergently transcribed from promoter regions of protein-coding genes (Quinn and Chang 2016; Grzechnik et al. 2014; Jensen et al. 2013). Whether the evolutionary conservation of this principle reflects functional importance of CUTs or their transcription, or simply that they are non-functional by-products of the basic mechanics of transcription, remains an open question.

The ~850 SUTs in budding yeast are detectable in proliferating wild-type cells, and are processed differently from CUTs (Neil et al. 2009; Tuck and Tollervey 2013). SUTs could be considered analogous to the previously annotated lncRNAs in fission yeast, which can be readily detected in proliferating wild-type cells (Wilhelm et al. 2008; Rhind et al. 2011) and show less variable expression than novel lncRNAs across the different conditions (*Figure 3A*). CUTs and SUTs almost exclusively originate from nucleosome-depleted regions at the ends of coding genes (Jensen et al. 2013). Our nucleosome-profiling data also suggest a strong tendency of Bidirectional and Intergenic lncRNAs to initiate in nucleosome-depleted regions upstream of positioned nucleosomes (*Figure 7*). These nucleosome data are only approximate, as we did not determine the profiles under the different genetic or physiological conditions when lncRNAs are more highly expressed.

The ~1700 budding yeast XUTs are derepressed upon deletion of the cytoplasmic exonuclease Xrn1 (ortholog of *S. pombe* Exo2), and are mostly antisense to mRNAs and anti-correlate with sense expression (van Dijk et al. 2011). These findings resemble our results (*Figure 4, Figure 5*). The targeting of XUTs by a cytoplasmic exonuclease implies their efficient export to the cytoplasm. However, the proposed inhibitory functions of XUTs on coding transcription are likely mediated co-transcriptionally, and so the relevance of their cytoplasmic export is unclear (Hansen et al. 2013; Tuck and Tollervey 2013). In budding yeast, XUTs are targeted by the NMD pathway before being degraded by Xrn1, and this pathway can be regarded as the last filter to dampen lncRNA expression (Malabat et al. 2015; Wery et al. 2016). Consistent with the cytoplasmic exonuclease acting as a backup system, we find that many CUTs and DUTs are also targeted by Exo2 and the NMD factor Upf1 (*Figure 2, Figure 4C*). Our XUTs and Antisense lncRNAs that engage with ribosomes, but not other classes of lncRNAs, are significantly enriched for RNAs derepressed in *upf1Δ* cells, supporting a role of the NMD in Exo2 degradation. Although there are overlaps of lncRNA expression in the absence of Upf1 and Exo2, much fewer lncRNAs are derepressed in *upf1* than in *exo2* mutants (*Figure 2, Figure 4C*). Absence of two other regulators of cytoplasmic RNA degradation, Ski7 and Pan2 (Lemay et al. 2010; Wolf and Passmore 2014), has only subtle effects on lncRNA derepression (*Figure 2, Figure 4C*). These results suggest that Exo2 plays the major role and can degrade lncRNAs independently of these other factors. However, a limitation of the current study is that RNA-seq only measures steady-state RNA levels, which integrates transcription and degradation. Findings from budding yeast indicate that mRNA levels can be adjusted by buffering mechanisms, allowing compensation of increased degradation by increased transcription or *vice versa* (Haimovich et al. 2013; Sun et al. 2017). Xrn1, the Exo2 ortholog, is required for this buffering. So it is possible that the weak derepression phenotypes of cytoplasmic RNA degradation mutants, other than *exo2*, reflect that lncRNA levels are efficiently buffered in these mutants. More work is required, however, to investigate whether the buffering system is conserved in fission yeast and whether lncRNAs are subject to it.

Budding yeast has no analogous lncRNA class to the DUTs defined here, because the RNAi pathway is missing (Harrison et al. 2009). Our results show that RNAi is important to control lncRNA expression in fission yeast, most notably Antisense lncRNAs that are derepressed in late meiosis (*Figure 5, Figure 6*). In fission yeast, RNAi can dampen RNA expression via either transcriptional or post-transcriptional mechanisms (Castel and Martienssen 2013; Smialowska et al. 2014). Our data do not allow to distinguish between these two possibilities. Although RNAi is not required for antisense-mediated transcriptional repression at three meiotic mRNAs (Chen et al. 2012), our and other results (Bitton et al. 2011) indicate a prominent global role of RNAi to suppress many Antisense lncRNAs. About 75% of all DUTs are Antisense lncRNAs. RNAi plays an even more important role than Exo2 in repressing Antisense lncRNAs, but also targets Bidirectional and Intergenic lncRNAs (*Figure 5A,B*). It is not clear whether the RNAi machinery is involved to a similar extent in controlling lncRNAs in multicellular organisms.

NUTs are another class of unstable lncRNAs in budding yeast, which substantially overlap with CUTs and XUTs (Schulz et al. 2013). NUTs are detected upon depletion of Nrd1, a member of the Nrd1-Nab3-Sen1 (NNS) complex that promotes transcriptional termination of lncRNAs (Schulz et al. 2013). NNS-mediated termination occurs in a TRAMP-dependent manner (Tudek et al. 2014). Nrd1 and Nab3 binding motifs are depleted in mRNAs but enriched in NUTs, indicating that NNS selectively terminates this class of lncRNAs. Fission yeast, however, does not show a similar motif bias (unpublished observation). Furthermore, no NNS complex was identified in fission yeast, and depletion of Seb1 impairs polyA-site selection but not RNA abundance (Lemay et al. 2016). Thus, a class corresponding to NUTs does not appear to exist in fission yeast. This conclusion is also consistent with Pab2, rather than TRAMP, being more important for exosome-mediated degradation of lncRNAs.

The different lncRNA classes based on RNA-processing pathways, while useful, are fairly arbitrary and overlapping. The lncRNAs are targeted by multiple redundant or coordinating pathways in an intricate backup system, although one pathway is often dominant for a given lncRNA. Moreover, RNA-processing mutants can lead to cellular re-routing of RNA degradation. Accordingly, there are substantial overlaps between different lncRNA classes in both budding and fission yeast. We find that cells require either the nuclear exosome or cytoplasmic exonuclease to survive, with the absence of both pathways being lethal. While these pathways function in other aspects of RNA metabolism, this synthetic lethality points to the importance of dampening the extensive lncRNA expression, which are much more affected in the corresponding mutants than are mRNAs (*Figure 2*). It is possible that the cytoplasmic exonuclease can serve as a backup to degrade transcripts that escaped degradation by the nuclear exosome. On the other hand, cells survive without the cytoplasmic exonuclease and RNAi or without the nuclear exosome and RNAi. Surprisingly, cells lacking both the cytoplasmic exonuclease and RNAi show fewer derepressed XUTs and DUTs than cells lacking only one of these pathways (*Figure 2*). This suppression might reflect that lncRNAs that cannot be degraded by RNAi are effectively targeted by the nuclear exosome. Consistent with this possibility, absence of both the nuclear exosome and RNAi leads to poor growth and large numbers of derepressed lncRNAs (*Figure 2, Figure 4D*). These findings indicate partially redundant roles for the nuclear exosome and RNAi pathways, which can back each other up with respect to many RNA targets. These two nuclear pathways can also degrade most XUTs that are further targeted by the cytoplasmic exonuclease. The RNAi and exosome pathways in fission yeast have overlapping functions to repress aberrant transcripts (Bühler et al. 2008; Zhang et al. 2011; Zofall et al. 2009) as well as meiotic mRNAs and other genomic regions (Yamanaka et al. 2013, 2010; Sugiyama and Sugioka-Sugiyama 2011). Our study highlights that the nuclear exosome and RNAi pathways also cooperate to suppress thousands of lncRNAs.

The expression levels of most lncRNAs are highly induced in non-dividing states (stationary phase and quiescence) and during meiotic differentiation, most notably in late meiosis when over 3000 Antisense lncRNAs are induced (*Figure 1, Figure 6*). These results raise the possibility that lncRNAs function during these conditions. It is known that unstable lncRNAs, normally targeted for rapid degradation, can become stabilised and functional under specialized conditions (Camblong et al. 2007; Houseley et al. 2008). Environmentally regulated changes to RNA quality-control activities can alter transcriptomes and mediate stress responses (Joh et al. 2016). RNA-processing pathways might become down-regulated under certain physiological conditions, allowing lncRNAs to accumulate. The mRNA levels of relevant RNA-processing genes do not strongly change in response to our physiological conditions, although mRNAs encoding nuclear-exosome components decrease ~2.7-fold during meiosis (Bitton et al. 2015a, and data not shown). Many meiotic mRNAs are repressed in mitotic cells by the RNAi and exosome pathways and derepressed during meiosis (Yamanaka et al. 2013, 2010; Sugiyama and Sugioka-Sugiyama 2011). Derepression of lncRNAs during meiosis and other specialized conditions could involve similar regulation. Indeed, our findings indicate that Exo2 also plays an important role in repressing many lncRNAs, but also many middle meiotic genes (Mata et al. 2002) that are derepressed in *exo2* mutants and meiosis. These results put Exo2 on the map as an important new regulator of meiotic gene expression.

In addition to derepression, the induction of lncRNAs could involve increased transcription (Castelnuovo et al. 2014). RNA-processing factors likely regulate RNA levels via coordinated interplays between transcription and degradation (Haimovich et al. 2013; Sun et al. 2017), and changes in this coordination could lead to the accumulation of different lncRNAs in different physiological conditions.

Antisense transcripts are the most widespread class of lncRNAs. In our data, over 70% of coding sequences produce at least one Antisense lncRNA from the other strand. This finding complements and extends previous analyses on antisense transcription in fission yeast (Wilhelm et al. 2008; Dutrow et al. 2008; Ni et al. 2010; Rhind et al. 2011; DeGennaro et al. 2013; Eser et al. 2016; Chen et al. 2012; Bitton et al. 2011; Zofall et al. 2009; Clément-Ziza et al. 2014; Zhang et al. 2011; Marguerat et al. 2012). Antisense lncRNAs include CUTs, DUTs, XUTs and other lncRNAs, with XUTs and especially DUTs being strongly enriched (*Figure 5A,B*). Thus, several RNA processing pathways can be involved in controlling Antisense lncRNAs. Previous studies have reported repressive effects of Antisense lncRNAs on their sense mRNAs (Bitton et al. 2011; Chen et al. 2012; Leong et al. 2014; Ni et al. 2010; Marguerat et al. 2012). Accordingly, we find a strong global tendency towards anti-correlation between Antisense lncRNA-mRNA expression levels under physiological conditions (*Figure 5C*). In contrast, Antisense lncRNA expression shows a slight tendency towards positive correlation with mRNA expression under the genetic conditions (*Figure 5C*). Thus, stabilization of Antisense lncRNAs in the absence of different RNA-processing factors appears generally not to be sufficient for mRNA repression. This finding suggests that Antisense lncRNAs often control mRNA expression at the level of transcription (e.g. by transcriptional interference or altered chromatin patterns) rather than functioning as transcripts. Alternatively, many Antisense lncRNAs might simply reflect opportunistic transcription, enabled by down-regulation of the dominant, overlapping mRNAs, with the anti-correlated expression reflecting passive, indirect effects. The ~29% of protein-coding regions not associated with Antisense lncRNAs are enriched for highly expressed genes, suggesting that these genes are either protected from, or interfere with, antisense transcription. Despite the global anti-correlation (*Figure 5C*), large numbers of Antisense lncRNAs go against this trend, indicating that the expression relationships between lncRNAs and mRNAs involve multiple processes and cannot be explained by a few regulatory or indirect mechanisms. This conclusion is consistent with the diverse findings on antisense lncRNA processes in other organisms (Pelechano and Steinmetz 2013; Mellor et al. 2016).

## Conclusions

This comprehensive study increases the number of lncRNAs annotated in fission yeast by almost 5-fold. The novel lncRNAs are typically lowly expressed but become highly derepressed in response to different genetic and physiological perturbations. In stark contrast, the mRNAs and annotated lncRNAs show less widespread changes in expression, especially in the genetic perturbations. The nuclear exosome, RNAi machinery, and cytoplasmic exonuclease are the dominant RNA-processing pathways degrading lncRNAs, used to define the CUTs, DUTs and XUTs, respectively. Bidirectional lncRNAs are enriched for CUTs and translating ribosomes, and positively correlate with divergent mRNA expression. Antisense lncRNAs are enriched for DUTs and XUTs, are mostly derepressed in late meiosis, and negatively correlate with sense mRNA expression in physiological, but not in genetic conditions. Intergenic lncRNAs are enriched for lncRNAs that are not classified as CUTs, DUTs or XUTs. The transcripton of Intergenic and Bidirectional lncRNAs initiates from regions that in wild-type cells are nucleosome-depleted, just upstream of a positioned nucleosome. Given their low expression and other features, it seems likely that any regulatory functions mediated by most lncRNAs are in *cis* and co-transcriptional.

Our findings highlight a substantial role of RNAi, in coordination with the nuclear exosome, in controlling a large number of lncRNAs typified by the new class of DUTs. Moreover, the findings reveal a prominent new function of the Exo2 cytoplasmic exonuclease, together with RNAi, in dampening the expression of both lncRNAs and mRNAs that become derepressed during meiosis. The nuclear exosome and cytoplasmic exonuclease together play an essential role for cell viability. The three RNA-processing pathways show overlapping roles and can target most lncRNAs with different affinities, forming an intricate, intertwined RNA-surveillance network. Besides these biological insights, this study provides broad data on diverse lncRNA characteristics and a rich resource for future studies on lncRNA functions in fission yeast and other organisms.

## Materials and methods

### S. pombe strains

All strains, physiological and growth conditions (Edinburgh minimal media, EMM2, or Yeast Extract media, YE), and biological repeats used for RNA-seq in the current study are detailed in ***Supplementary files 1 and 2.*** Biological repeats are based on independent cell cultures of the different strains. The number of different samples was decided to broadly interrogate key genetic and physiological conditions, balancing biological insight with costs. The PCR-based approach (Bähler et al. 1998) was used for gene deletions of *exo2, pan2*, and strains used to generate double mutants. Double mutants among *dcr1*, *rrp6* and *exo2* were created by crossing the corresponding single mutants (***Supplementary file 1***). Strain *h^−^dcr1::nat ura4^−^*was generated using the PCR-based approach (Bähler et al. 1998). Random spore analysis was used to create the other single mutant strains with the correct mating-types. Strains were crossed and incubated on malt extract agar (MEA) for 2-3 days at 25°C. Tetrads were treated with zymolyase (0.5 mg/ml, MP Biomedicals Europe) and incubated at 37°C for at least 4 h to release spores. Spores were germinated on YE agar plates before being replica plated to selective EMM2 plates as appropriate. All deletion junctions were PCR verified (Bähler et al. 1998). Crosses and selection by random spore analysis were as follows: *h^+^ ade6-M216 leu1-32 ura4-D18 his3-D1 rrp6::ura4* was crossed with *h^−^ura4-D18* with selection on EMM2 plates to create *h^+^ rrp6::ura4 ura4-D18; h^−^exo2::kanMX6 ade6-216* was crossed with *h^+^ ura4-D18* with selection on YE + kanamycin plates, and on EMM2 plates with or without uracil, to select for *h^+^ exo2:: kanMX6 ura-D18* and *h^−^exo2:: kanMX6 ura-D18*. Tetrad analysis was used to analyse the meiotic products resulting from the crosses in all combinations of the *rrp6Δ, dcr1Δ* and *exo2Δ* single mutant strains. Strains were crossed and incubated on MEA plates for 2-3 days at 25°C. The resulting tetrads were dissected using a micromanipulator (Singer Instruments), and spores germinated on YE plates after 5 days of growth. Haploid colonies arising from germinated spores were then streaked to selective plates to test for KAN, NAT and URA markers. All deletion junctions in double-resistant colonies were PCR-verified (Bähler et al. 1998).

### Growth conditions

All mutant cell cultures were harvested at mid-log phase (optical density, OD595 = 0.5). For stationary-phase experiments, wild-type cells were grown in EMM2 at 32°C. A sample representing 100% survival was harvested when cultures reached a stable maximal density. Colony forming units (CFUs) were measured every 24 hours after this initial time-point, and another sample harvested when cultures reached 50% survival. For quiescence experiments, cells were grown in EMM2 at 32°C an OD600 of 0.2, before being centrifuged, washed twice in EMM2 without nitrogen (NH4Cl) and cultured in EMM2 without nitrogen at 32°C. Cells under nitrogen starvation reached an OD600 of 0.8 within 24 hours, and were harvested at 24 hours and 7 days after nitrogen removal. For meiotic timecourses, *pat1-114* diploid cells were grown to mid-log phase before being shifted to EMM2 without nitrogen. Cells were incubated at 25°C overnight to synchronise them in G1 phase. Meiosis was induced by addition of NH4Cl to final concentration of 0.5 g/L and incubation at 34°C (0 hour timepoint). Cells were harvested by centrifugation of 50 ml cultures at 2300 rpm for 3 min, and pellets were snap-frozen and stored at -80°C prior to RNA extraction.

### RNA-seq experiments and initial analyses

RNA was extracted from harvested cells using a hot-phenol method (Bähler and Wise 2017). The quality of total extracted RNA was assessed on a Bioanalyser instrument (Agilent). Strand-specific RNA-seq libraries were prepared using an early version of the Illumina TruSeq Small RNA Sample Prep Kit. For polyA-enriched samples, library preparation and sequencing protocols were as described by Lemieux et al. (2011), and for samples depleted for rRNAs (*rrp6Δ, exo2Δ*), as described (Bitton et al. 2014). RNA-seq libraries were sequenced on an Illumina HiSeq 2000 instrument, using single-end runs with 50 bp reads (The Berlin Institute for Medical Systems Biology, Germany). Reads were aligned to the fission yeast genome with the exonerate software (Slater and Birney 2005), and reads matching to multiple genomic locations were assigned at random to one of these locations. Reads containing up to 5 mismatches (not clustered at read ends) were kept for further analysis. Between 20 and 50 million mappable reads were obtained for each library (~80-85% of total reads were mappable). Expression scores were calculated for annotated features using the genome annotation available in PomBase on 9th May 2011 (Wood et al. 2012). Reads per kilobase of transcript per million reads mapped (RPKMs) for annotated features correlated strongly between biological replicates (*rPearson* >0.98). Mapping and expression score pipelines were performed as described (Lemieux et al. 2011).

### Segmentation of sequence data to define novel lncRNAs

A simple heuristic was designed to detect novel lncRNAs from RNA-seq data. This segmentation heuristic was optimised for its ability to detect the 1557 annotated lncRNAs, and validated by visual inspection of RNA-seq data. Custom scripts for segmentation of RNA-seq data were written in *R* and *Perl*. The following RNA-seq data from initial sequencing runs were pooled (2 biological repeats each): *rrp6-ts, exo2Δ, dis3-54, pab2Δ, nmt1-mtr4* (Lemay et al. 2014; Bitton et al. 2015a), *ago1Δ, rdp1Δ, dcr1Δ, pan2Δ, upf1Δ*, Stat 100%, Stat 50%, Quies 24 h, Quies 7d, Meiotic pool (Schlackow et al. 2013), YE1, and Refence (control) (***Supplementary files 1 and 2***). Segments were delimited from the pooled data using a 10 hits/bp cut-off. Segments <100 bp apart and differing in pooled read density (average hits/bp of segment) by <10-fold were joined together. Using the PomBase genome annotation (May 2011), segments overlapping annotations on the same strand, including untranslated regions, were removed. We discarded segments of <200 bp as lncRNAs are defined by an arbitrary minimal length cut-off of 200 nt (Mattick and Rinn 2015), reflecting RNA-seq library protocols that exclude small RNAs. The remaining consecutive segments >200 bp defined 5775 novel lncRNAs.

### Analyzing of RNA expression

Novel lncRNAs defined by the segmentation process described above, together with all annotated transcripts, were analysed using the Bioconductor DESeq2 package (Love et al. 2014). Differentially expressed genes were defined as those being >2-fold induced (average of two biological repeats) or repressed and showing significant changes (adjusted p <0.05) compared to three reference samples as determined by DESeq2. For hierarchical clustering of expression data, log2 ratios were clustered in R with the pheatmap package, using the Euclidian distance measure and the ward.D or ward.D2 clustering options. For expression correlation analyses, we evaluated the similarity of expression levels of Bidirectional and Antisense lncRNAs to the expression levels of their neighbouring mRNAs using normalised expression values across the the entire dataset. Vectors of mRNA-lncRNA pairs were generated and Pearson’s correlation coefficients were computed. For lncRNAs associated with multiple mRNAs, only the nearest mRNA was considered. Functional enrichments of gene lists were performed using the AnGeLi tool which applies a 2-tailed Fisher’s exact test (Bitton et al. 2015b).

### Classification into CUTs, DUTs and XUTs

Differential expression data from *dcr1Δ*, *exo2Δ* or *rrp6Δ* mutants were filtered to retain only transcripts that were significantly induced in ≥1 mutant compared to wild-type controls (expression ratio >2 and adjusted p-value < 0.05). RPKM values from independent biological repeats for the differentially expressed RNAs were then standardized to have a mean value of 0 and a standard deviation of 1, followed by clustering using the Mfuzz clustering function in R (Kumar and E. Futschik 2007). The number of clusters “c” was set to 3 and the fuzzification parameter “m” to 1.25. To further reduce ambiguity when associating RNAs to clusters, the minimum membership value of a lncRNA belonging to a specific cluster was set to 0.7 (Kumar and E. Futschik 2007). For this classification, more recent PomBase annotations were used which contained only 1533 annotated lncRNAs (7308 annotated and novel lncRNAs in total).

### Classification into Bidirectional, Antisense and Intergenic lncRNAs

To assess whether a given annotated or novel lncRNA overlaps with any mRNAs in either orientation, we systematically aligned the lncRNA coordinates relative to the annotation in Ensembl *S. pombe*, Assembly ASM294v2, release 33 (Flicek et al. 2014), enhanced by a modified annotation set that better delineates transcript boundaries. To this end, we exploited Transcription Start Sites (TSS) determined using Cap Analysis of Gene Expression (CAGE) (Li et al. 2015) and Transcription Termination Sites (TTS) defined using genome-wide polyadenylation site mapping (Mata 2013). For genes without these higher quality boundaries, we used the annotated TSS and TTS. All TSS and TTS used are provided in ***Supplementary file 7***. We called overlaps in either orientation when ≥1 nt was shared between transcripts. Using the same criteria, we tested for overlaps with novel lncRNAs that have been recently reported (Eser et al. 2016) (***Supplementary file 4***).

Using these overlap criteria, we classified all known and novel lncRNAs based on their proximity to nearby mRNAs. We defined lncRNAs as Intergenic if they do not overlap with any nearby mRNA, Sense-overlapping lncRNAs if they overlapped with any mRNA on the same strand, Antisense if they overlap ≥1 nt with a mRNA on the opposite strand, and Bidirectional if their TSS was <300 nt up- or down-stream of a TSS of a mRNA on the opposite strand. Naturally, given the compact fission yeast genome, there was some overlap between these classes. We reassigned lncRNAs present in two classes using the following criteria. The lncRNAs classified as both Bidirectional and Intergenic (482 lncRNAs) or as Bidirectional and Antisense (1068 lncRNAs) were assigned to Bidirectional lncRNAs only. The 135 lncRNAs classified as both Sense-overlapping and either 86 Antisense or 49 Bidirectional lncRNAs were, respectively, assigned to Antisense or Bidirectional lncRNAs only.

### Translation analysis of lncRNAs

For the ribosome profiling analysis, we systematically looked for overlaps between the translated regions defined by Duncan & Mata (2014) and all annotated and novel lncRNAs. The significance of enrichments among different lncRNA classes was determined using the *prop.test* function in R.

### Growth phenotypes of mutant cells

For a semi-quantitative analysis of cell growth, we used spot assays. After overnight pre-culture, yeast cells were adjusted to the same OD value (~0.6). For each strain, 5 µl of six serial (five-fold) dilutions were spotted onto YE plates and grown at 32°C.

### Nucleosome profiling

Mononucleosomal DNA (MNase digested) from exponentially growing wild-type (*972 h^−^*) cells in EMM2 was generated as reported (Lantermann et al. 2009). Two independent biological repeats were performed. Sequencing libraries from MNase-digested DNA were prepared using the NEBNext ChIP-Seq Library Prep Master Mix Set for Illumina (E6240S). Pair-end 50 bp reads were obtained with an Illumina MiSeq sequencer at the Genomics and Genome Engineering Facility at the UCL Cancer Institute. MNAse sequencing data was mapped using Bowtie2 (Langmead and Salzberg 2012). Nucleosome maps for visualisation were performed with nucwave (Quintales et al. 2015), following the web recommendations (http://nucleosome.usal.es/nucwave/). Data were analysed using the ‘*computeMatrix reference-point*’ and ‘*heatmapper’* functions from the deeptools package with transcription start sites as reference points (Ramírez et al. 2014).

### Data availability

Sequencing data have been submitted to ArrayExpress and the European Nucleotide Archive under accession numbers PRJEB7403, E-MTAB-708, E-MTAB-2237, MTAB-1154, E-MTAB-1824 (RNA-seq; for sample accessions, see ***Supplementary files 1and 2***) and PRJEB21376 (nucleosome profiling).

## Acknowledgements

We thank Philipp Korber for comments on the manuscript, Wei Chen (Berlin Institute for Medical Systems Biology) for help with RNA-sequencing, and Pawan Dhami (Genomics and Genome Engineering Facility at UCL Cancer Institute) for help with sequencing for the nucleosome profiling. This work was funded by a Wellcome Trust Senior Investigator Award (grant number 095598/Z/11/Z) and a Royal Society Wolfson Research Merit Award to JB. SM was supported by the Medical Research Council, FB by a grant from the Canadian Institutes of Health Research (MOP-106595), and JM by a BBSRC project grant [BB/M021483/1].

## Additional files

### Supplementary files

Supplementary file 1. RNA metabolism mutants used.

Supplementary file 2. Physiological conditions used.

Supplementary file 3. Expression values for all RNAs and all conditions.

Supplementary file 4. Overlap of novel lncRNAs with those reported by Eser et al. (2016).

Supplementary file 5. ncRNA classes and information on overlaps between lncRNAs and coding sequences or mRNAs.

Supplementary file 6. Polysome profiling analysis of actively translated lncRNAs.

Supplementary file 7. All TSS and TTS used.

Supplementary file 8. lncRNA regions showing unusually high histone occupancy.

## Figure supplements

Figure 1-figure supplement 1. Comparison of RPKM values for samples depleted for rRNAs vs samples enriched for poly(A) RNAs.

Figure 6-figure supplement 1. Expression changes of CUTs, DUTs and XUTs in different physiological conditions.

---

**Figure 1 – figure supplement 1.**
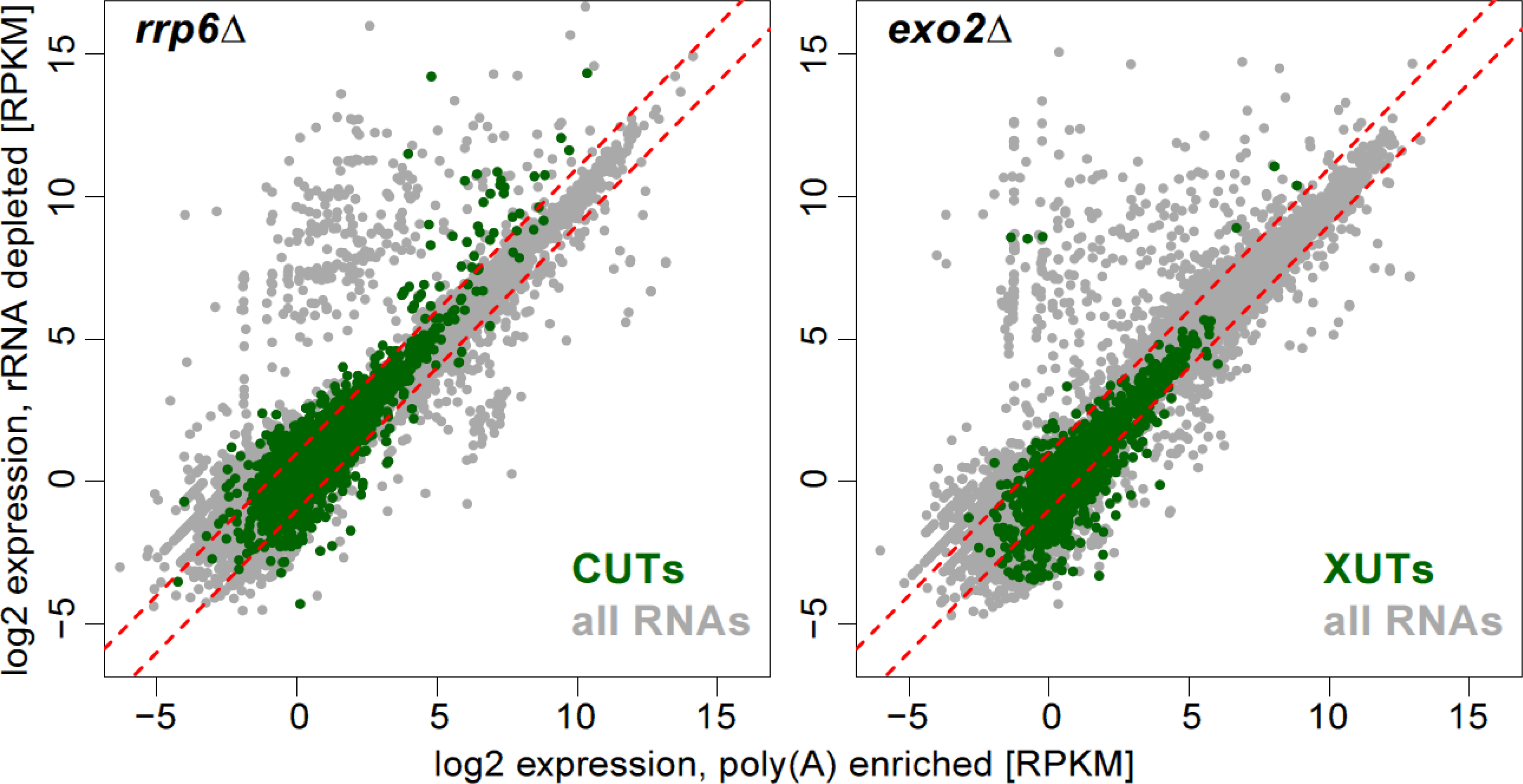
Comparison of RPKM values for samples depleted for rRNAs vs samples enriched for poly(A) RNAs. Samples were prepared from *rrp6Δ* (left) or *exo2Δ* (right) mutants, with CUTs and XUTs, respectively, highlighted in green and all other coding and non-coding RNAs shown in grey. Red dotted lines: 2-fold changes. Only 325 and 354 RNAs were induced in rRNA depleted *rrp6Δ* and *exo2Δ* samples, respectively, compared to the corresponding poly(A)-enriched samples. In both cases, 242 of those RNAs are known to lack poly(A) tails, including tRNAs, snRNAs, snoRNAs and mitochondrially-encoded RNAs. Only 49 and 23 novel lncRNAs were induced in rRNA depleted *rrp6Δ* and *exo2Δ* samples, respectively.

**Figure 6 - figure supplement 1.**
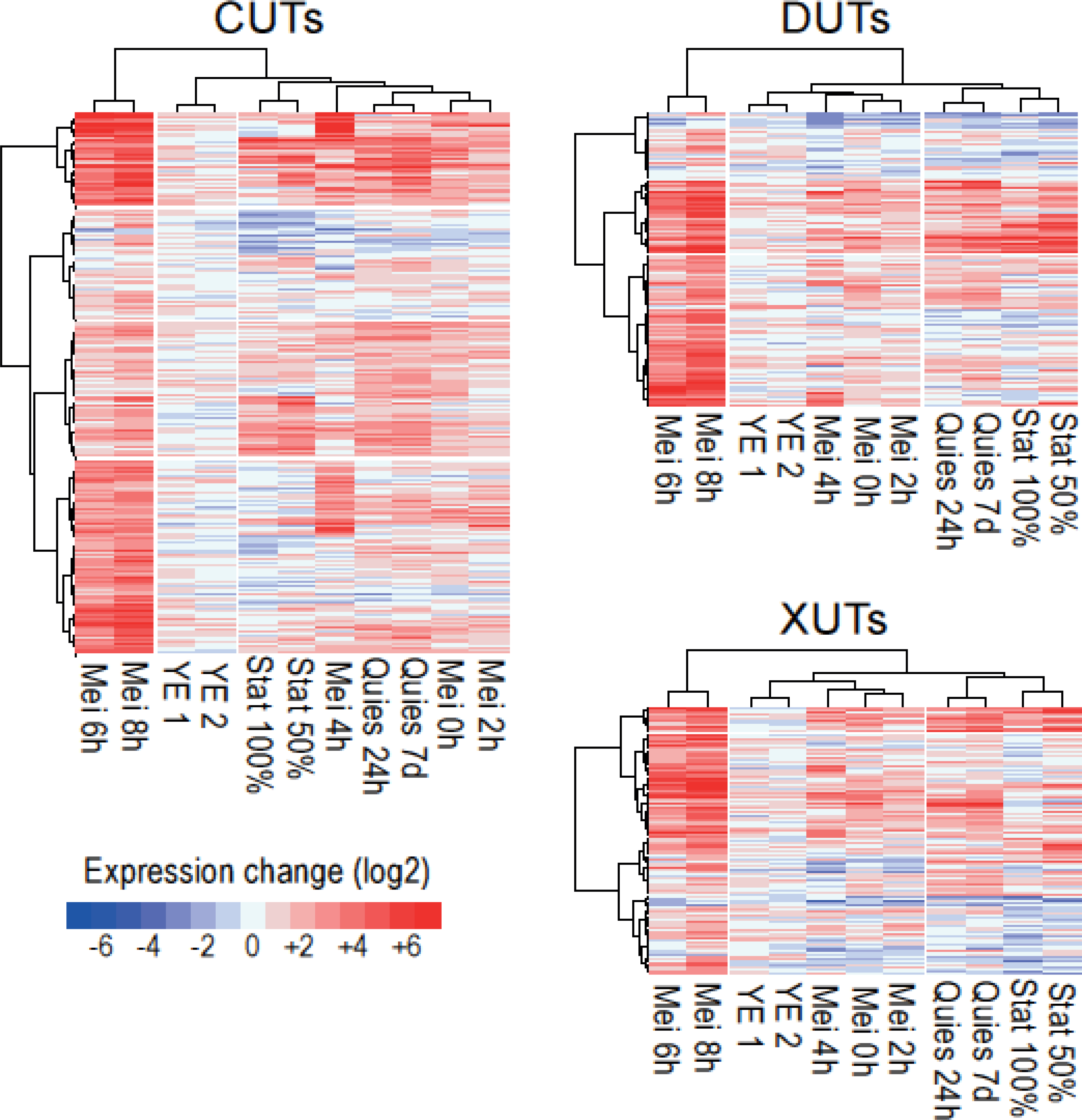
Expression changes of CUTs, DUTs and XUTs in different physiological conditions. Hierarchical clustering of physiological conditions for all CUTs, DUTs and XUTs as indicated. The R pheatmap package was used for clustering applying the Euclidean distance and the ward.D clustering option. Changes in RNA levels in response to the different physiological conditions (indicated at bottom) relative to control cells grown in minimal medium are color-coded as shown in the colour-legend at bottom right (log2 fold-changes). The physiological conditions cluster into three groups: late meiosis (Mei 6-8h), stationary phase (Stat 50-100%), and quiescence/early meiosis (Quies 24h/7d, Mei0-4h).

